# Reduced accrual of mineral-associated organic matter after two years of enhanced rock weathering in cropland soils, though no net losses of soil organic carbon

**DOI:** 10.1101/2024.06.23.600278

**Authors:** Noah W. Sokol, Jaeeun Sohng, Kimber Moreland, Eric Slessarev, Heath Goertzen, Radomir Schmidt, Sandipan Samaddar, Iris Holzer, Maya Almaraz, Emily Geoghegan, Benjamin Houlton, Isabel Montañez, Jennifer Pett-Ridge, Kate Scow

**Affiliations:** Lawrence Livermore National Laboratory, Physical & Life Science Directorate, Livermore, California, USA; University of California, Davis, Department of Land, Air and Water Resources, Davis, California, USA; United States Department of Agriculture, California Climate Hub, Davis, CA, USA; Yale University, Department of Ecology and Evolutionary Biology, New Haven, CT, USA; University of California, Davis, Institute of the Environment, Davis, California, USA; University of California, Santa Barbara, Environmental Studies Program, Santa Barbara, California, USA; Yale Center for Natural Carbon Capture, Yale University, New Haven, CT, USA; Cornell University, Department of Ecology and Evolutionary Biology & Department of Global Development, Ithaca, New York, USA; Dept. of Earth and Planetary Sciences, Univ. of California, Davis, CA, 95616; University of California Merced, Life & Environmental Sciences Department, Merced, CA, USA; Innovative Genomics Institute, Berkeley CA, USA

## Abstract

Enhanced rock weathering (ERW), the application of crushed silicate rock to soil, can remove atmospheric carbon dioxide by converting it to (bi)carbonate ions or solid carbonate minerals. However, few field studies have empirically evaluated ERW in field settings. A critical question remains as to whether additions of crushed rock might positively or negatively affect soil organic matter (SOM) – Earth’s largest terrestrial organic carbon (C) pool and a massive reservoir of organic nitrogen (N). Here, in three irrigated cropland sites in California, USA, we investigated the effect of crushed meta-basalt rock additions on different pools of soil organic carbon and nitrogen (i.e., mineral-associated organic matter and particulate organic matter), active microbial biomass, and microbial community composition. After two years of crushed rock additions, mineral-associated organic matter (MAOM) stocks were lower in the upper surface soil (0-10 cm) compared to unamended controls. At the two sites where baseline pre-treatment data were available, neither total SOC nor SON decreased over the two years of study in crushed rock or unamended control plots. However, the accrual rate of MAOM-C and MAOM-N at 0-10 cm was lower in plots with crushed rock vs. unamended controls. Before ERW is deployed at large scales, our results suggest that field trials should assess the effects of crushed rock on SOM pools, especially over multi-year time scales, to accurately assess changes in net C and understand the mechanisms driving interactions between ERW and SOM cycling.

## Introduction

To keep planetary warming below 2*°*C this century, the global economy must be rapidly decarbonized by reducing greenhouse gas emissions (Shukla et al. 2022). To meet this target, most Integrated Assessment Models also require direct removal of carbon dioxide (CO_2_) from the atmosphere – both to remove excess CO_2_ that has accumulated in the atmosphere and to offset emissions that cannot be abated (Fuss et al. 2018; Minx et al. 2018). Enhanced rock weathering (ERW) in soils has recently attracted much interest as a CO_2_ removal approach due to its relatively low cost, scalability, and potential co-benefits – especially on croplands (Kantola et al. 2017, 2023; Beerling et al. 2018). ERW accelerates the natural process of silicate weathering by adding finely-crushed silicate rock to soils, which can then react with dissolved atmospheric CO_2_, generating (bi)carbonate ions or carbonate minerals (Renforth 2012). As crushed silicate rock dissolves, base cations like the magnesium cation (Mg^2+^) and calcium cation (Ca^2+^) are released, increasing net alkalinity in the surrounding soil pore water (Gillman 1980; Holzer et al. 2023b). This increase in alkalinity ultimately removes CO_2_ from the atmosphere by increasing the amount of dissolved inorganic carbon (C) stored in groundwater, rivers, and the ocean. Some inorganic C may also persist as solid carbonates in soils, subsurface geologic deposits or sediments (Köhler et al. 2010; Guo et al. 2023). Modeling studies suggest that ERW could remove 0.5–2 billion tons of atmospheric CO_2_ per year if deployed on suitable croplands at a global scale (Beerling et al. 2020). At the same time, nutrients released from weathering crushed rock may also boost soil fertility and crop yields (Haque et al. 2019; Kelland et al. 2020) and could help displace some need for agricultural liming (Dietzen et al. 2018). However, while the theoretical potential of ERW is dramatic, few field studies have verified modeled estimates of CO_2_ removal or quantified how crushed rock interacts with biogeochemical cycles in working agricultural soil systems (Moosdorf et al. 2014; Goll et al. 2021; Vicca et al. 2022).

The effect of crushed rock additions on soil organic matter (SOM) pools is one critical unknown for quantifying the net C removed via ERW (Vicca et al. 2022). Soil organic matter is Earth’s largest terrestrial C pool and a key source of nitrogen (N) and other essential nutrients that fuel ecosystem productivity (Scharlemann et al. 2014). While increases to SOM with crushed rock could enhance the C removal benefits of ERW by drawing down more atmospheric CO_2_, reductions in SOM could diminish or negate its C drawdown benefits. Understanding the effects of crushed rock on SOM stocks is a critical gap in the measurement, reporting, and verification of C removed through ERW (Holzer et al. 2023a; Suhrhoff et al. 2024). Moreover, potential positive or negative effects of crushed rock on soil organic nitrogen (SON) has key implications for soil fertility and crop productivity in agroecosystems (Kantola et al. 2023). Amidst rapidly growing interest and investment in ERW from federal governments and voluntary C markets, it is vital that measurement, reporting, and verification protocols for ERW are informed by field studies that account for the full range of effects of crushed rock on the soil system.

Enhanced rock weathering may lead to SOM accrual or loss through a suite of interconnected pathways. Weathered silicate rocks not only release base cations, but also generate reactive secondary minerals. These reactive secondary minerals are the primary location where organic matter (OM) is stored in agricultural soils – known as ‘mineral-associated organic matter,’ or MAOM (Cambardella and Elliott 1992; Sokol et al. 2022b). On average, ∼80% of SOM in global croplands is MAOM; the remaining portion exists as particulate organic matter, or POM (Sokol et al. 2022b). Newly created reactive mineral sites that are produced from weathered silicate rock could increase the amount of MAOM in soil by providing more surface sites for OM to bind (Slessarev et al. 2021). Rock weathering can also release Ca^2+^ ions, which can enhance SOM accrual via sorption of OM on mineral surfaces via cation bridging, co-precipitation with carbonate minerals, and production of microbial biofilms involved in MAOM formation (Rowley et al. 2018; Shabtai et al. 2023; Buss et al. 2024). Crushed rock amendments could also shift soil biogeochemical processes in ways that cause net OM mineralization (Yan et al. 2023), potentially leading to decreased SOM stocks or reduced rates of SOM accrual. For instance, an increase in pH or in the release of micro-nutrients under ERW can induce higher microbial activity, leading to decomposition of SOM (Fang et al. 2023). At the same time, if the addition of crushed rock leads to increased production of organic acids by plant roots or soil microbes to chemically weather the rock and gain access to nutrients, these organic acids could destabilize mineral-organic associations and lead to SOM loss (Keiluweit et al. 2015). The emergence of positive or negative ERW-SOM effects will likely be influenced by the activity and functional capacity of the soil microbial community, since soil microorganisms are key agents in the formation and decomposition of SOM – particularly MAOM (Kallenbach et al. 2015; Sokol et al. 2022a, 2024). In addition to their impacts on SOM cycling, soil microbes are intimately associated with rock weathering, often residing on the surfaces of soil minerals and accelerating mineral dissolution via biological weathering (Uroz et al. 2009).

Here, in a series of ERW field trials on three different croplands in the Central Valley of California, USA, we examined how two years of meta-basalt crushed rock amendments affected SOM pools and soil microbial communities. We assessed the impacts of ERW treatments on SOC and SON stocks, MAOM and POM pools, active soil microbial biomass, and microbial community composition at two depth increments (0-10 cm and 10-30 cm). We asked: (1) What are impacts of ERW on SOC and SON stocks and on different SOM pools (MAOM vs. POM)? (2) What is the effect of ERW on active soil microbial biomass and bacterial and fungal community composition? (3) How do these effects vary across the three field trials, each with different soil properties, cropping systems, and management approaches?

## METHODS

### Experimental Design and Soil Sampling

Research was conducted at three ERW field trials on different croplands in the Central Valley of California, USA, all initiated in fall 2019 and managed by the Working Lands Innovation Center, administered by the University of California, Davis, Institute of the Environment (Fig. S1, Table S1). The ‘Yolo’ ERW field trial was located at the UC Davis Campbell Tract Agricultural Research Station in Davis, Yolo County, California (38° 31’53.96’N, 121°46’54.15’W; mean annual temperature = 15.6 °C; mean annual precipitation = 449 mm; data obtained from California Irrigation Management Information System station #6, 1983-2022). The Yolo trial was on a field annually cropped with corn (*Zea mays* L.) in 2020 and 2021, following the start of the ERW field trial in 2019. The other ERW field trials (‘Merced-1’ and ‘Merced-2’) were located on two different fields on a private farm in Los Banos, Merced County, California (37° 7’10.099’N, 120°44’38.41’W; MAT =16.7 °C, MAP = 174 mm; data obtained from California Irrigation Management Information System station #124, 1996-2022). The Merced-1 site was on a field continuously cropped with alfalfa (*Medicago sativa* L.) since the start of the field trial in fall 2019. The Merced-2 trial was on a field with an annual crop rotation of corn (2019), tomato (2020; *Solanum lycopersicum* L.), and cilantro (2021; *Coriandrum sativum* L.). Soils at the Yolo site are coarse-loamy, mixed, superactive, nonacid, thermic Mollic Xerofluvents. Soils at Merced-1 are fine-loamy, mixed (calcareous), thermic Typic Haplaquolls; and soils at Merced-2 are fine-loamy, montmorillonitic (calcareous), thermic Typic Epiaqualfs. The Yolo site was irrigated via subsurface drip, while Merced-1 and Merced-1 were irrigated via furrow/flood irrigation. Further site details – including soil properties and site management information – are described in Table S1. The Yolo site has also previously been described in Holzer et al. (2023b).

Meta-basalt crushed rock was sourced from Specialty Granules LLC, from the Ione mine in Amador County, California (Ione, CA, USA). The rock is derived from Late Jurassic volcanic units that include meta-basalt and meta-andesite with minor dacite and rhyolite (Gutierrez et al. 2015; Holland 2016). The median grain size of crushed rock was 102-107 µm; elemental composition was determined using lithium borate fusion followed by inductively coupled plasma optical emission spectroscopy (ICP-OES; Bureau Veritas Mineral Laboratories, Richmond, BC, Canada; Table S2). In fall 2019 and fall 2020, 40 t ha^-1^ of crushed rock were applied to experimental plots (*n* = 3 at the Yolo site; *n* = 5 at the Merced sites). Following application, crushed rock was tilled in at all three sites; paired control plots (without rock amendments) were also tilled and managed in the same way as plots with crushed rock. Merced-1 received a rock application only in fall 2019 since alfalfa (a perennial) was grown on this field. At the Yolo site, plots with crushed rock were 0.045 ha in size, control plots were 0.09 ha (Fig. S1). At the Merced sites, plots with crushed rock and control plots were both 0.6 ha in size. In total, there were 26 plots across the three field trials: *n* = 13 plots with crushed rock and *n* = 13 control plots (Fig. S1; Table S1).

In November/December 2021, soil cores were collected at approximately equal intervals along the length of each plot using handheld augers, separated into 0-10 cm and 10-30 cm depth increments, and then thoroughly homogenized to generate one composite sample for each plot × depth increment. Soil cores were collected at least 30 feet away from the edge of the field and 6 feet from adjacent plots. Five cores were collected per plot at Yolo, and 10 cores were collected per plot at Merced (due to differences in plot size). From this composite sample, a ∼10 g subsample was stored at -80°C in a Whirl-Pak bag for subsequent DNA extraction. A separate ∼100 g subsample of fresh soil was immediately brought back to the lab for gravimetric soil water content, water holding capacity, active microbial biomass, and cumulative respiration measurements (described below). The remaining soil was air-dried until reaching constant moisture, then passed through a 2-mm sieve for further edaphic analyses. In December 2021, a Geoprobe (Geoprobe Systems, Saline, KS, USA) was deployed at the sites to determine plot-specific bulk density at 0-10 and 10-30 cm depths.

### Soil Analyses

Soil pH and electrical conductivity (EC) were measured using a benchtop pH meter and 10-g samples of air-dried soil mixed 1:2 with deionized water. Gravimetric soil moisture was determined as the mass difference between a fresh soil sample and a soil sample dried at 105°C for 48 hours. To determine water holding capacity (WHC), fresh soils were wet to 100% field capacity by saturating a subsample of soil and then allowing each sample to drain for 2 hours. Water holding capacity was determined as the mass difference between wet soils and soils dried at 105°C for 48 hours (Strickland et al. 2019).

Soil organic matter (SOM) was separated via a physical fractionation assay into three fractions: (1) dissolved organic matter (DOM), (2) mineral-associated organic matter (MAOM), and (3) particulate organic matter (POM; Bradford et al. 2008). First, 10-g of air-dried soil was weighed into a 50-mL falcon tube; 40-mL of deionized water was added to the tube and the soil solution was vortexed and placed on a shaker table for 15 min at 200 rpm. After shaking, samples were centrifuged for 15 min at 3400 rpm; the supernatant was then filtered through a 0.45-µm syringe filter (the < 0.45-µm extract was defined as DOM). To separate the POM and MAOM fractions, 30-mL of 5% sodium hexametaphosphate (NaHMP) was added to the soil pellet at the base of each 50-mL falcon tube. Samples were vortexed and placed on a shaker table for 16 h at 200 rpm. After shaking, the soil solution was passed through a 53-µm sieve. The <53-µm fraction (clay + fine silt fraction; defined as MAOM) and the >53-µm fraction (sand, coarse silt, and large organic matter fragments; defined as POM) were oven-dried to constant mass at 105°C. Dried samples were finely ground to a powder-like consistency with a mortar and pestle.

An acid fumigation method was used to differentiate between organic and inorganic C in the solid-phase MAOM and POM fractions (Brodie et al., 2011). Two separate aliquots of each dried and ground MAOM and POM sample were prepared for elemental analysis. The first set of aliquots (‘non-acid fumigated’) were immediately weighed into tin capsules. The second set of aliquots (‘acid-fumigated’) were weighed into glass 20-mL scintillation vials, then placed in a vacuum-sealed glass desiccator containing a glass beaker with 50-mL of 12 mol L^-1^ hydrochloric acid. Samples were fumigated for 48 h; the duration was determined by analyzing a subset of soil samples at 12, 24, 48, and 96 h of fumigation, and selecting the minimum time in which all carbonates were removed (Komada et al. 2008). Following acid-fumigation, samples were dried at 60°C for 16 h, cooled at room temperature in a desiccator cabinet, stirred with a glass rod to return the sample to a fine powder, and weighed into tin capsules.

All samples were analyzed for elemental (% C, % N) and isotope composition (δ^13^C, δ^15^N), on an elemental analyzer coupled to an isotope ratio mass spectrometer (EA-IRMS). Elemental and isotopic analyses were conducted at the Yale Analytical and Stable Isotope Center (New Haven, Connecticut, USA). The amount of inorganic C was calculated as the difference in % C between the non-acid fumigated (i.e., soil containing organic and inorganic C) and the acid-fumigated aliquot (i.e., soil containing organic C only) in each MAOM and POM sample. The acid-fumigated aliquots of MAOM and POM were used for organic C, δ^13^C, N and δ^15^N measurements. We present SOM data in two different ways:

1. *SOM stocks at the final timepoint (Fall 2021):* Stocks of C and N in MAOM, POM, and total SOM (MAOM + POM) in fall 2021 were calculated using the concentrations of C and N in each fraction, the plot-specific bulk density value (corrected for the volume of crushed rock per g of soil, as well as the mass of coarse fragments), and the thickness of the sample layers (i.e., 10-cm or 20-cm), so that they were expressed in g C or N m^-2^ at a given depth increment (i.e., 0-10 cm or 10-30 cm; Sokol et al. 2017). Plot-specific bulk density values for 2021 were corrected for the added mass of rock in soil by subtracting the mass of crushed rock estimated to be in each gram of soil, adjusted for the proportion of crushed rock estimated to be in the 0-10 cm increment and the 10-30 cm increment. At the Yolo and Merced-2 sites, 50% of added crushed rock was assumed to be in the 0-10 cm depth increment and 50% in the 10-30 cm depth increment, based on the depth of tillage at these sites. At Merced-1, all crushed rock was assumed to be in the 0-10 cm depth increment. On average, crushed rock accounted for a small portion (< 3%) of the total soil volume.
2. *Plot-specific two-year change in SOM concentrations:* At two of the three sites (Yolo and Merced-2), pre-treatment baseline soil samples were collected in each plot in fall 2019, before the experimental period began. We performed the same SOM fractionation procedure described above on these samples to determine plot-specific baseline concentrations of SOM pools for all experimental plots (bulk density values were not collected at this time point, so it was not possible to calculate stocks). We used these pre-treatment baseline measurements to calculate plot-specific changes through time in SOM concentrations (MAOM, POM, and total SOM) at Yolo and Merced-2 over the two-year period, by subtracting the pre-treatment baseline value (i.e., in fall 2019) from the final value (i.e., in fall 2021).

Baseline data at Yolo indicated that control plots had higher initial SOM concentrations than plots where crushed rock was subsequently applied (Table S3). At Merced-2, there were no significant differences in initial SOM concentrations between control plots and plots where crushed rock was subsequently applied (Table S3). These initial differences at Yolo were explicitly accounted for when measuring plot-specific changes in SOM concentrations through time, as described above. However, these differences may have biased differences in SOM stock values at Yolo in 2021, by leading to an overestimation of the effect of crushed rock on SOM pools. To account for these initial differences at Yolo when calculating differences in SOM stocks in 2021 between rock and control plots, we included a correction factor when measuring the % difference between SOM stocks in rock versus control plots in 2021. The corrected % differences for each SOM pool at Yolo in 2021 was calculated as:

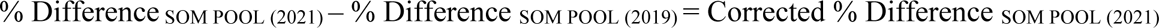

where ‘% Difference _SOM POOL (2021)_’ is the average difference in a given SOM pool between plots with crushed rock versus control plots, and % Difference _SOM POOL (2019)_ is the average baseline difference between these plots, before treatments were applied (Table S5). In the Results section below, we present both the uncorrected and corrected % difference for SOM stocks at Yolo in 2021. While no baseline data was collected at Merced-1, the experimental design at Merced-1 was similar as Merced-2 (i.e., staggered treatments plots, in contrast with Yolo where control plots were clustered toward one end of a field; see Fig. S1). This experimental design minimized differences in baseline values between treatment plots at Merced-2; we therefore assumed that differences in pre-treatment baseline values at Merced-1 were also not significantly different.

### Dissolved organic matter and elemental abundances

Dissolved organic matter (from samples extracted with deionized water, as described above) was analyzed on a TOC Analyzer for total C and on a GasBench (Thermo Delta Plus XP) for δ^13^C at the Yale Analytical and Stable Isotope Center. The DOM extract was also analyzed for elemental abundances of 7 elements: magnesium (Mg), calcium (Ca), sodium (Na), silicon (Si), aluminum (Al), iron (Fe), and phosphorous (P) using inductively-coupled plasma mass spectrometry (ICP-MS), on ThermoScientific XR High-Resolution ICP-MS Lawrence Livermore National Laboratory (Livermore, CA, US).

### Microbial Analyses

Active microbial biomass was estimated using the substrate-induced respiration (SIR) method on 10-g of fresh soil (Fierer et al. 2003). Autolyzed yeast extract solution (12 g L^−1^) was added to the soil sample in a sealed jar with a rubber septum (2:5 soil:liquid ratio by mass), and the CO_2_ respiration rate was measured from 0.5 to 3 h using an infrared gas analyzer (LiCOR Biosciences; Li850; Slessarev et al. 2020). Cumulative CO_2_ respiration was measured on 10-g of fresh soil, adjusted to 65% WHC, and incubated in a sealed mason jar for 60 days at 20°C. The cumulative CO_2_ production was measured using an infrared gas analyzer (LiCOR Biosciences; Li850).

We extracted DNA from frozen soil samples using a Qiagen DNEasy PowerSoil Pro kit (following the manufacturer’s instructions) on three separate 0.5-g aliquots per sample, which were then combined into a single composite replicate. Microbial community profiling was performed on an Illumina MiSeq 2 as paired end for 2 x 250 cycles, at Lawrence Livermore National Laboratory (Livermore, CA, USA). The V4 region of 16S rRNA gene was targeted for bacteria and the ITS2 region was targeted for fungi on an Illumina Miseq platform. The V4 region of 16S rRNA gene was amplified using primers 515F (5′-GTGYCAGCMGCCGCGGTAA -3′) and 806R (5′-GGACTACNVGGGTWTCTAAT-3′) and the ITS2 region of fungi were amplified using the primers 5.8S-Fun (5′-AACTTTYRRCAAYGGATCWCT-3′) and ITS4R (5′ - AGCCTCCGCTTATTGATATGCTTAART-3′; Apprill et al. 2015; Parada et al. 2016; Taylor et al. 2016). Once raw sequences were obtained, ‘cutadapt’ (Martin 2011) was used to remove primers and ‘dada2’ was used for further processing of sequences (Callahan et al. 2016). For analysis, 16S rRNA gene forward reads were truncated to 250 base pairs (bp) and reverse reads to 200bp based on quality profile plots. Reads were truncated when the quality score was below 30. The fungal ITS sequences did not have their forward and reverse reads truncated to a specific length since the ITS region is known to be highly variable in length (Feibelman et al. 1994; Schoch et al. 2014); trimming and filtering were conducted using the same parameters as 16S. After quality processing, paired sequences were dereplicated and the ‘dada’ function was used to denoise the unique reads based on the error rates calculated with the ‘learnErrors’ function (Callahan et al. 2016). Once paired sequences were merged and chimera were removed, the SILVA database (Quast et al. 2013) and UNITE database (Nilsson et al. 2019) were used to assign taxonomy to 16S and ITS sequences, respectively. All 16S rRNA sequences that were assigned to chloroplast, eukaryota, and mitochondria were removed prior to statistical analyses.

### Statistics

Statistical analyses were conducted in R (R Core Team, 2023). The effect of crushed rock on SOM fractions (both SOM stocks in 2021 and two-year change in SOM concentrations), active microbial biomass, cumulative respiration, and other soil properties were calculated with linear mixed effect models using the *lme4* package (Bates et al. 2015). For all models, treatment (rock versus control) and site (Yolo, Merced-1, Merced-2) were included as fixed effects; plot was included as a random effect. Non-significant interaction terms were dropped from the model. Models for each depth increment (0-10 cm and 10-30 cm) were fit separately (Slessarev et al. 2020). All models were screened for normality of residuals (Shapiro-Wilk test) and heteroscedasticity of residuals (visual assessment of residual plots). For microbial community data (16S and ITS), differences in bacterial and fungal community (Bray-Curtis and Euclidean) were assessed through permutational multivariate analysis of variance (Permanova). Beta diversity was visualized by principal coordinates analysis (PCoA) based on Bray-Curtis dissimilarity. Differential abundance analysis of bacterial and fungal ASVs was conducted using the “DeSeq2” package (McMurdie and Holmes 2013; Love et al. 2014). For all models, p < 0.05 was considered significant, and p < 0.1 was considered marginally significant.

## RESULTS

### Soil Organic Matter Pools

After two years of crushed meta-basalt applications, total SOC and SON stocks were lower in the surface soil (0-10 cm) of plots amended with crushed rock relative to paired control plots (p = 0.02 and p = 0.007, respectively; Table S4). There was no effect of crushed rock on total SOC or SON at 10-30 cm (p = 0.2 and p = 0.7, respectively, Table S4). The effect of crushed rock on surface soil SOC and SON stocks was driven by lower mineral-associated organic matter (MAOM), which comprised a majority of the SOM pool at the three field sites (MAOM was ∼65% of total SOM; the remaining ∼35% was POM). At the end of the two-year experimental period (fall 2021), crushed rock was associated with lower surface soil MAOM-C stocks by 159 ± 37 g C m^-2^ (p = 0.001; Table S4; Fig. 1). On average, MAOM-C stocks were lower in plots with crushed rock by 16 ± 5 % at Merced-1, by 11 ± 6 % at Merced-2, and by 13 ± 12 % at Yolo (or by 30 ± 12 % at Yolo when no correction factor was applied, see Methods above; Table S5). In fall 2021, crushed rock was associated with lower surface soil MAOM-N stocks by 17 ± 5 g N m^-2^ (p = 0.001; Table S4). On average, MAOM-N stocks were lower in plots with crushed rock by 14 ± 4 % at Merced-1, by 11 ± 8 % at Merced-2, and by 13 ± 6 % at Yolo (or 27 ± 4 % at Yolo when no correction factor was applied, Table S5).

**Figure 1.**
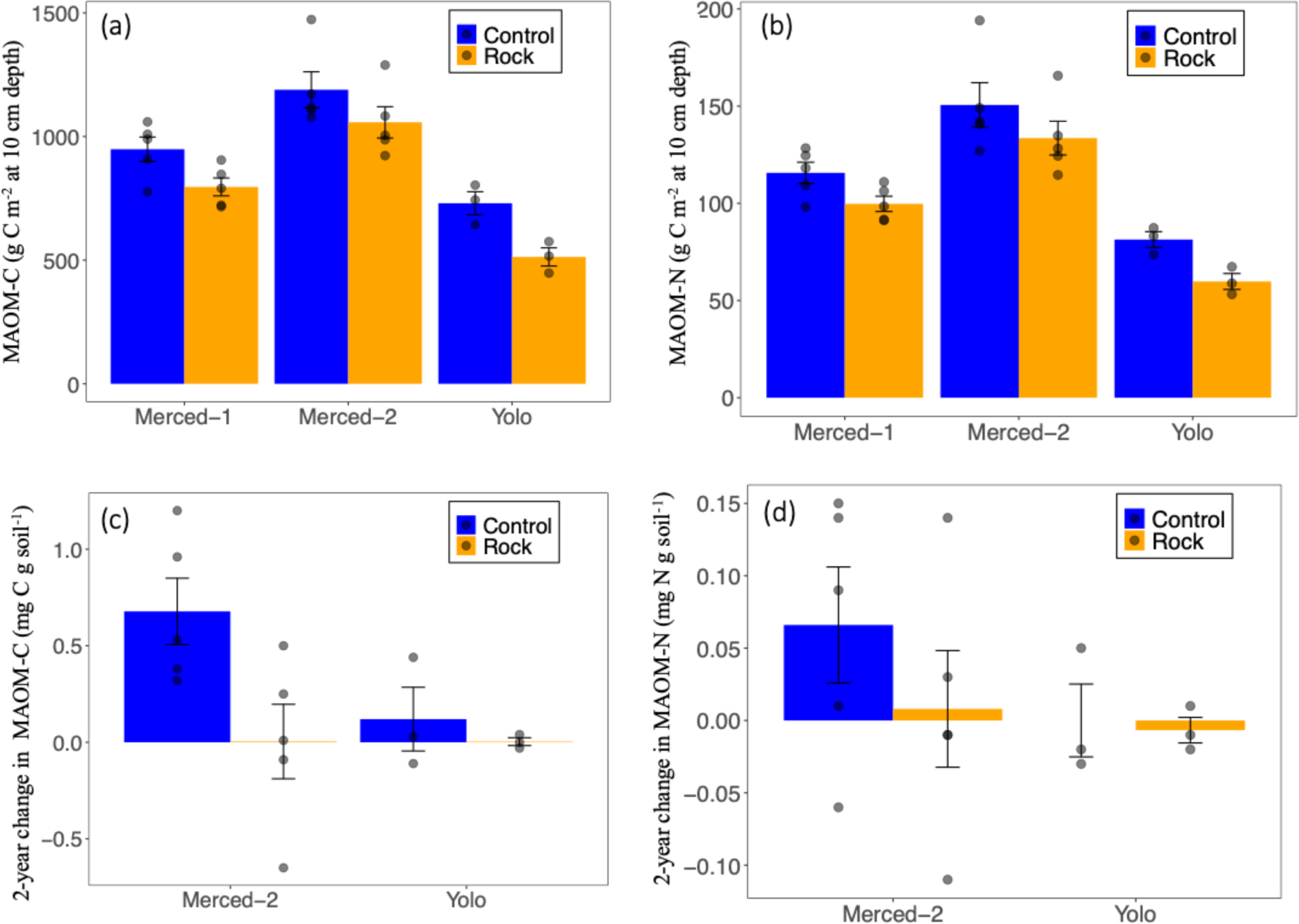
Mineral-associated organic matter C and N stocks in upper surface soil (0-10 cm) of plots with crushed meta-basalt rock versus unamended control plots across three cropland enhanced rock weathering field trials in California, USA. Stocks of (a) mineral-associated organic matter C (MAOM-C), and (b) mineral-associated organic matter N (MAOM-N) at 0-10 cm depth in plots amended with crushed rock (orange bars) relative to control plots (blue bars) in fall 2021. Stocks expressed as g C (or N) m^-2^ by thickness of soil layer (i.e., 10 cm). There were 13 replicates (*n = 13*) per treatment across all field trials. At the sites where baseline data was available (Yolo and Merced-2), we calculated plot-specific changes in the concentration of (c) MAOM-C (mg C g soil^-1^) and (d) MAOM-N (mg N g soil^-1^) over the two-year period. There were 8 replicates (*n* = 8) across two field trials with baseline data. Changes in MAOM-C or N over time were measured by change in concentration (mg C or N g soil^-1^) in fall 2021 versus pre-treatment baseline values in fall 2019.

On average, there were no net losses of total SOM, SON, MAOM or POM fractions over the two-year experimental period in both crushed rock and control plots. At Yolo and Merced-2 (the two sites where pre-treatment baseline data was available), the two-year changes in MAOM-C and MAOM-N concentrations were net positive in crushed rock and control plots (Fig.1c,d; Table S7). However, the two-year change in MAOM-C concentration (i.e., the accrual rate) was lower in plots with crushed rock versus control plots, and this effect varied by site (site*treatment interaction, p = 0.07; Table S6). At Merced-2, the accrual rate of MAOM-C was lower in plots with crushed rock by an average of 0.67 ± 0.26 mg C g soil^-1^ over the two-year period, while at Yolo, the accrual rate of MAOM-C was lower in plots with crushed rock by an average of 0.12 ± 0.17 mg C g soil^-1^ (Table S7).

There was no significant effect of crushed rock at 0-10 or 10-30 cm depths on POM-C or POM-N stocks in fall 2021(Table S4). Similarly, there was no significant effect of crushed rock on the two-year change in POM-C or POM-N concentrations at 0-10 or 10-30 depth (Table S6). While the POM pool did not exhibit a consistent, significant response to crushed rock, it did trend either toward a positive response in the presence of crushed rock, or a less negative response than the MAOM fraction (Table S5, S7). The contrasting responses of the POM and MAOM pools influenced how the total SOM pool responded to crushed rock amendments. At all three sites, responses of total SOC and SON stocks were more muted than MAOM-C and MAOM-N pools (Table S5). At Merced-1, for example, MAOM-C stocks were 16 ± 5% lower in plots with crushed rock compared to control plots, whereas total SOC stocks were only 9 ± 4 % lower in plots with crushed rock compared to control plots (Table S5). Moreover, while the two-year change in MAOM-C and MAOM-N concentrations were significantly lower in plots with crushed rock relative to control plots at 0-10 cm depth (Fig. 1c, Table S6), there was no significant difference between crushed rock and control plots in total SOC and SON concentrations at 0-10 cm depth (p = 0.5; Table S5).

There was no effect of crushed rock on the size of the DOC pool at 0-10 cm or 10-30 cm depth (Table S3), though the δ^13^C value of the DOC fraction was more enriched in δ ^13^C in plots with crushed rock relative to control plots at 10-30 depth (p < 0.05; Table 1). There was no effect of crushed rock on the δ^13^C or δ^15^N values of the MAOM or POM fraction at either 0-10 cm or 10-30 cm depth (Table S7, S8).

**Table 1.** Modeled regression coefficients of the effect of site and treatment (crushed rock) on several soil variables in three temperate cropland enhanced rock weathering (ERW) field trials. Shown below are coefficient estimates (± standard error, SE) from linear mixed effect models where crushed rock had a significant effect on: (a) soil pH at 0-10 cm, (b) gravimetric soil water content at 0-10 cm (% soil moisture), (c) soil water holding capacity at 10-30 cm (% soil moisture), (d) δ^13^C of DOC at 10-30 cm. There were 13 replicates (*n = 13*) per treatment across all field trials*. (p < 0.1 is considered marginally significant and indicated by a single asterisks *; p < 0.05 is considered significant and indicated by two asterisks **)*.

| | Coefficient estimate ± SE | p value |
| --- | --- | --- |
| **(a) pH at 0-10 cm** |  |  |
| Intercept | 7.88 ± 0.06 | <0.0001** |
| Site (Merced-2) | -0.26 ± 0.06 | 0.0002** |
| Site (Yolo) | -0.84 ± 0.07 | <0.0001** |
| Treatment (crushed rock) | 0.1 ± 0.05 | 0.06* |
| **(b) Gravimetric Soil Water Content at 0-10 cm (% soil moisture)** |  |  |
| Intercept | 0.2 ± 0.005 | <0.0001** |
| Site (Merced-2) | 0.03 ± 0.006 | 0.0003** |
| Site (Yolo) | -0.15 ± 0.007 | <0.0001** |
| Treatment (crushed rock) | -0.01 ± 0.006 | 0.07* |
| **(c) Water Holding Capacity at 10-30 cm (% soil moisture)** |  |  |
| Intercept | 30.56 ± 0.7 | <0.0001** |
| Site (Merced-2) | -3.92 ± 0.76 | <0.0001** |
| Site (Yolo) | -2.41 ± 0.9 | 0.01** |
| Treatment (crushed rock) | -1.3 ± 0.7 | 0.06 |
| **(d) $\delta^{13}\text{C}$-DOC at 10-30 cm** |  |  |
| Intercept | -25.30 ± 0.41 | <0.0001** |
| Site (Merced-2) | 0.71 ± 0.50 | 0.17 |
| Site (Yolo) | 1.67 ± 0.58 | 0.008** |
| Treatment (crushed rock) | 1.48 ± 0.44 | 0.003** |

### Soil inorganic carbon and soil properties

At Yolo and Merced-2 (the two sites with pre-treatment baseline data), the two-year change through time in soil inorganic C (SIC) concentration was greater in plots with crushed rock relative to control plots at 0-10 cm (Fig 2f; Table S6). In fall 2021, there was no effect of crushed rock on SIC stocks at 0-10 cm, though SIC stocks trended higher in plots with crushed rock at 10-30 cm (p = 0.1; Fig 2c). Crushed rock was associated with higher pH in the surface soil (0-10 cm) relative to control plots by 0.1 ± 0.05 units (p = 0.09; Table 1). At 0-10 cm, gravimetric soil water content was lower in plots with crushed rock relative to control plots in fall 2021; at 10-30 cm, water holding capacity was lower in plots with crushed rock relative to control plots (Table 1). There was no effect of crushed rock on electrical conductivity (EC) at either depth, or on total elemental abundances (Mg, Ca, Si, Na, Al, Si, P) in the DOM extract from bulk soil (Table S10).

**Figure 2.**
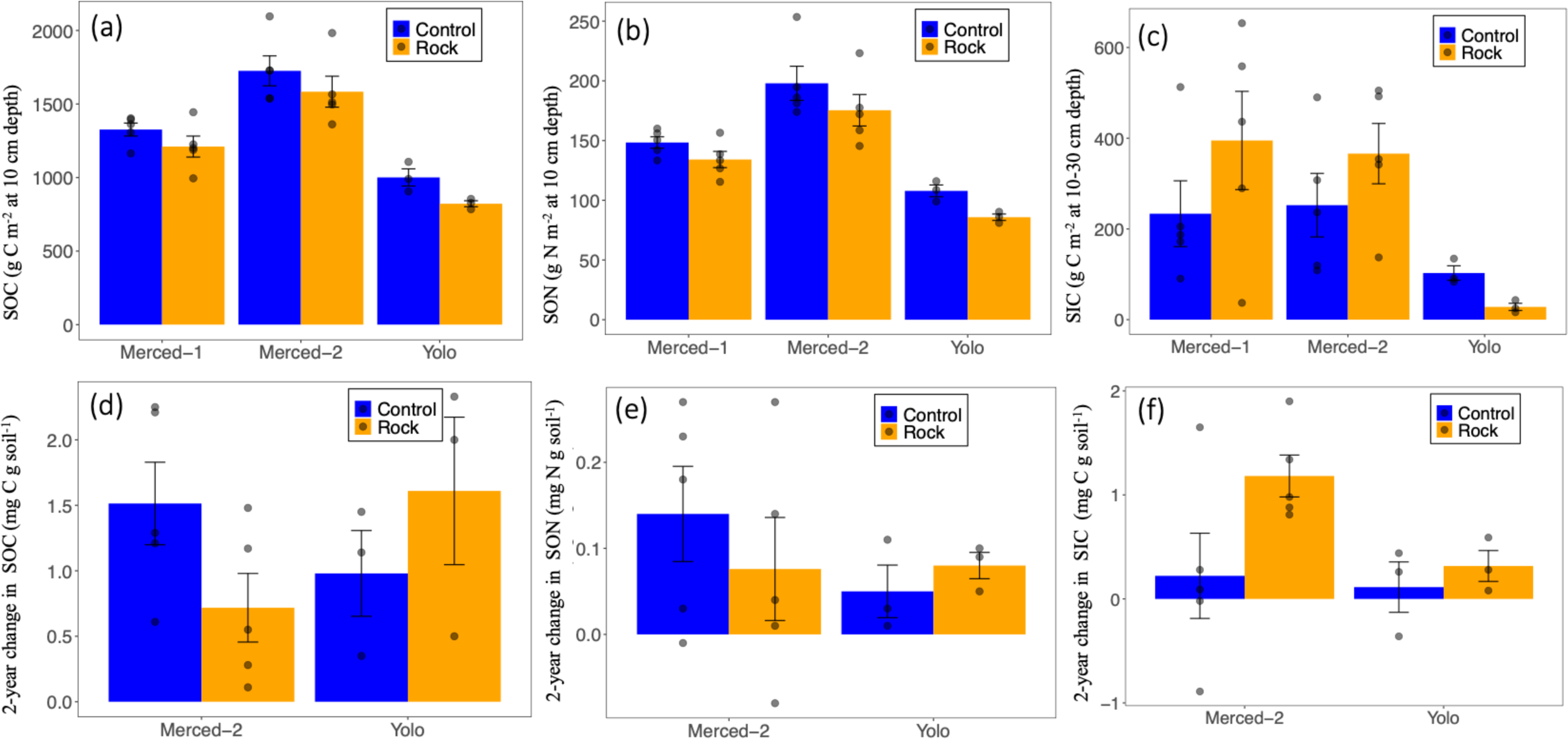
Total soil organic C and N and soil inorganic C in plots amended with crushed rock versus unamended control plots across three cropland enhanced rock weathering field trials in California, USA. Stocks of total (a) soil organic carbon (SOC) and (b) soil organic N (SON) at 0-10 cm depth, and (c) stocks of soil inorganic C at 10-30 cm depth measured in fall 2021 in plots with crushed rock versus control plots. At the two sites where baseline data was available (Yolo and Merced-2), we calculated the plot-specific change through time in the concentrations of (d) SOC, (e) SON, and (f) SIC over the two-year period. Changes in SOM pools over two years were measured as the change in concentration (mg C or N g soil^-1^) in fall 2021 compared to pre-treatment baseline values in fall 2019. There were 13 replicates (*n = 13*) per treatment across all field trials, and 8 replicates (*n* = 8) across two field trials with baseline data. Stocks are expressed as g C or N per m^-2^ by thickness of soil layer (10 cm or 20 cm); concentrations are expressed in mg C or N g soil^-1^.

### Microbial biomass and community composition

Active microbial biomass in surface soil was lower in plots with crushed rock than control plots (Fig. 3a; Table S10). There was a strong positive linear relationship between active microbial biomass and MAOM-C stocks in surface soil in fall 2021 across all three sites (Fig 3b, r^2^ = 0.74, p < 0.005). In contrast, active microbial biomass at 10-30 cm was greater in crushed rock than control plots (Table S10). There was no significant effect of crushed rock on cumulative CO_2_ efflux over a 60-d incubation period, measured on fresh soil harvested in Fall 2021 at 0-10 cm or 10-30 cm (Table S10).

**Figure 3.**
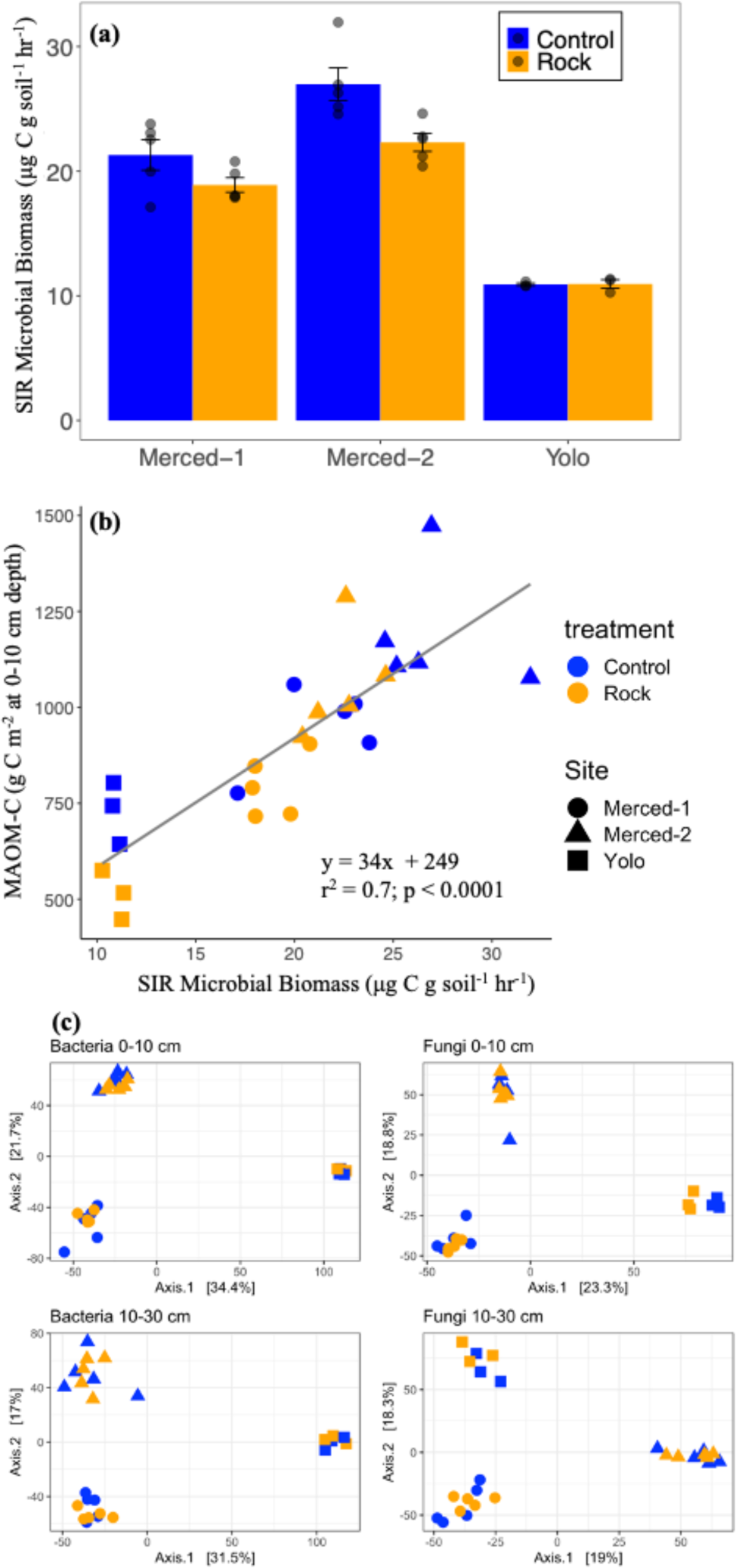
Active microbial biomass and community composition in upper surface soil (0-10 cm) of plots with crushed meta-basalt rock versus control plots across three cropland enhanced rock weathering field trials in California, USA. (a) Active microbial biomass (measured via substrate-induced respiration) at 0-10 cm depth in plots with crushed rock relative to control plots. b) Linear relationship between active microbial biomass and stocks of mineral-associated organic matter (MAOM-C) at 0-10 cm depth. (c) Principal coordinate analysis (PCoA) plots showing the compositional differences (via Euclidean distances) of ASVs for bacteria and fungi at 0-10 cm and 10-30 cm, respectively. There were 13 replicates (*n = 13*) per treatment across all field trials.

Despite differences in bacterial and fungal community composition among the three sites and two depths in fall 2021, there was no effect of crushed rock on microbial community composition (Fig. 3c, Table S11). Differential abundance testing indicated that a small number of bacterial and fungal taxa were more abundant under crushed rock relative to control plots (Table S12). At 10-30 cm, three alphaproteobacterial taxa in the family *Sphingomonadaceae*, and a strain of *Actinobacteria* classified in class *Thermoleophilia*, were more abundant in plots with crushed rock relative to control plots (Table S12). At 0-10 cm, two fungal taxa – *Microscales* and *Pezizales* (both in Ascomycota) – were more abundant in plots with crushed rock (Table S12).

## DISCUSSION

Before enhanced rock weathering is deployed at large scales in cropland soils, it is critical to characterize the full suite of its impacts on the ecosystem C balance (Goll et al. 2021). Here, we examined ERW effects on soil organic matter (SOM) pools. We demonstrated that after two years of meta-basalt crushed rock additions across three ERW cropland field trials in the Central Valley of California, total SOC and SON stocks were lower in the surface soil (0-10 cm) of plots amended with crushed rock relative to control plots (Table S4). This effect was the result of lower mineral-associated organic matter (MAOM) – the largest pool of OM in these soils (∼65% of total SOM), and typically the slower-cycling pool of SOM (Heckman et al. 2022). At the end of the two-year experimental period (i.e., in fall 2021), crushed rock was associated with lower surface soil MAOM-C stocks and MAOM-N stocks relative to unamended control plots by 159 ± 37 g C m^-2^ (p = 0.0001; Table S4) and 17 ± 5 g N m^-2^ (p = 0.001; Table S4). We did not find evidence that lower surface soil MAOM-C stocks led to a net loss of SOC or SON; the two-year change in plot-specific SOC and SON concentrations were net positive at the two sites where pre-treatment baseline data was available (Fig. 2d). However, the accrual rate of MAOM was lower in plots with crushed rock relative to control plots over the two-year period (Fig. 1; Table S7). Below, we discuss the effects of ERW on soil inorganic and organic C pools, how the microbial community may help explain the effects of crushed rock on SOM, and identify key next steps to better understand the interactions between rock weathering and OM cycling.

The two-year change in soil inorganic C (SIC) concentrations was higher in sites with crushed rock amendments compared to unamended controls by 0.68 ± 0.23 mg C g soil^-1^ (p = 0.017; Fig. 2f, Table S6). Since the cations released from weathered rock can form pedogenic carbonates under relatively high pH conditions (Huang et al. 2024), increased soil inorganic C can indicate effective ERW (Haque et al. 2020a). In particular, ERW should primarily yield solid phase carbonates at sites with calcareous soils, where soil water may be near saturation with respect to the excess cations released during weathering (Haque et al. 2020a). Indeed, SIC stocks and accrual rates were most pronounced in the calcareous soils of the Merced sites (Fig 2c, f; Table S6-S7). Similar increases in SIC have been observed in some agricultural soils under ERW: for example, an increase in SIC of ∼1.83 g CO_2_ kg soil^-1^ was observed over a two-year period in a cropland in Ontario, Canada under wollastonite application (Haque et al. 2020b). However, increases to solid phase carbonates are certainly not always observed under ERW, such as below a threshold pH value (Haque et al. 2019; Guo et al. 2023)

Surface soil SOM pools showed contrasting responses to additions of crushed rock. While MAOM stocks were consistently lower in plots with crushed rock than control plots at 0-10 cm (Fig. 1a), POM stocks at 0-10 cm depth trended higher in plots with crushed rock in surface soil (particularly at the Merced-1 and Yolo sites; Table S5). The contrasting responses of the MAOM and POM pools dampened the response of the total SOM pool to ERW (Fig. 2). For instance, while MAOM-C stocks ranged between 11 ± 6% to 16 ± 5% lower in plots with crushed rock additions, total SOC stocks ranged between 4 ± 6 and 9 ± 4 % lower in the presence of crushed rock. Moreover, while the two-year change in MAOM-C concentration (i.e., the accrual rate) was lower in plots with crushed rock by an average of 0.5 ± 0.2 mg C g soil^-1^ (p = 0.03; Table S4), there was no significant difference in the accrual rate of total SOC between control plots and plots with crushed rock (p = 0.5; Table S6). To understand how and why the POM pool may dampen the effect of ERW on total SOM, it will be critical to monitor the responses of the MAOM and POM pools over longer time scales, and to assess consequences for total SOM.

Overall, the effects we observed of crushed rock on SOM pools are roughly commensurate with how other agricultural management approaches – such as liming – may affect SOC pools over similar timescales. In our plots, we found that surface soil SOC stocks were lower in plots with crushed rock by an average of 140 g C m^-2^ at 10 cm depth (p = 0.02; Table S4). After three years of lime incorporation in an Australian Oxic Paleustalf, surface soil SOC stocks were ∼3.6 t C ha^-1^ (360 g C m^-2^) lower than un-limed controls,, or a ∼13% reduction in SOC stocks (Chan and Heenan 1999). After three lime applications in a Brazilian Planosol, SOC in the top 15-cm was reduced by 9-15% (Kowalenko and Ihnat 2013). Importantly, as we discuss further below, the magnitude and the direction of effects of liming on SOC can change over longer time-scales and ultimately become positive in some contexts (Paradelo et al. 2015). Thus, it will be critical to capture longer-term effects of ERW on SOM pools.

The different responses of surface soil MAOM and POM pools to ERW over the two-year period may be partially explained by the activities of the soil microbial community. Soil microorganisms are key agents of SOM accrual and decomposition; in particular with formation of MAOM (Kallenbach et al. 2015; Sokol et al. 2024). Given that active soil microbial biomass was significantly lower in the surface soil of plots with crushed rock (Fig. 3a) – and that active microbial biomass and MAOM-C stocks were strongly positively correlated across the three field trials (Fig. 3b) – lower active microbial biomass may help explain reduced MAOM accrual in these plots. One key microbial pathway for MAOM formation is decomposition of POM into smaller biomolecules that subsequently associate with mineral surfaces (Liang et al. 2017). One possibility is that the lower active microbial biomass associated with ERW translated into decreased microbial-mediated transformation of POM to MAOM. As a result, there was less formation of MAOM and greater accumulation of POM in the presence of crushed rock. In support of this hypothesis, we found that POM-C stocks and accrual rates often trended higher in crushed rock than control plots across sites (though this effect was not significant; Table S4-S8). At Yolo – the site with the lowest active microbial biomass (Fig. 3a) – we observed the greatest accrual of POM and the lowest accrual of MAOM (Fig. 1; Table S7). Moreover, we found no significant changes in SOM pools at 10-30 cm depth, where active microbial biomass was substantially lower than in the 0-10 cm increment. It is unclear if lower active microbial biomass in the surface soil of plots with crushed rock was a function of physicochemical changes induced to the soil, such as altered mobility and availability of nutrients with shifts in pH (Min et al. 2021). Alternatively, ERW may have reduced SOM by non-microbial mechanisms, and lower SOM led to reduced active microbial biomass, since microbial biomass and SOM tend to be strongly correlated (Liang et al. 2024). It will be important for future mechanistic studies to untangle the directions of these effects.

Other microbial mechanisms may explain decreased MAOM under ERW. While active microbial biomass was lower under ERW, it is possible that a subset of the microbial community exhibited greater microbial activity in the presence of crushed rock, due to changes in the physical and chemical environment. Heightened microbial activity under ERW could stimulate decomposition of a portion of the MAOM pool, leading to decreased net MAOM accrual rates in crushed rock than control plots. For instance, soil pH was higher in the surface soil of plots with crushed rock by 0.1 ± 0.05 units (p = 0.06; Table 1), and prior soil-water measurements conducted at the Yolo site demonstrated higher bicarbonate alkalinity in plots with crushed rock relative to control plots (Holzer et al. 2023b). Increased pH and alkalinity may stimulate microbial activity through several means, such as alleviating acid retardation of microbial growth and SOM consumption (Malik et al. 2018). As one parallel, agricultural liming has been linked with enhanced microbial activity and microbial priming of SOM stocks over shorter time scales (Paradelo et al. 2015). Short-term enhancement of C mineralization rate has been also reported in recent mesocosm studies on ERW (Yan et al. 2023). In support of enhanced microbial activity under ERW, we found a more enriched δ^13^C value of DOC in plots with crushed rock at 10-30 cm (Table 1), which can indicate greater microbial activity and priming of SOM (Krüger et al. 2023).

One key question is whether certain microbial taxa may exhibit greater activity under ERW, as microbial populations colonize fresh mineral surfaces, and simultaneously decompose OM to gain access to nutrients. If ERW selects for certain taxa that chemically weather rock but also destabilize organo-mineral interactions, these taxa may lead to decreased MAOM accrual. Differential abundance analysis on amplicon sequence data (16S and ITS) suggested that specific taxa were more abundant in the presence of crushed rock, even while community composition did not significantly differ between the two treatments (Fig 3c). Bacterial taxa in the *Sphingomonadaceae* and *Thermoleophilia* families were more abundant in the presence of crushed rock (Table S12); these families are known to play roles in mineral weathering (Huang et al., 2014; Wang et al., 2017; Varliero et al., 2021). Additionally, fungi in the genera *Microscales* and *Iodophanus* were more abundant in plots with crushed rock (Table S12). *Iodophanus* has been reported to positively respond to Ca^2+^ ions (Diorio, 1999). Such data can help guide future efforts to more directly target specific microbial taxa involved in ERW and SOM cycling, using methods like quantitative stable isotope probing (qSIP; Hungate et al., 2015).

Our findings raise several critical questions and identify mechanisms for future research to explore. Overall, our results suggest that ERW may reduce the accrual rates of MAOM in irrigated semi-arid croplands, necessitating studies that disentangle the various factors that influence how crushed rock may impact SOM pools. These factors include crop type, soil properties, management strategies like irrigation and cover cropping, climate, and the composition and amount of crushed rock, as well as application frequency and incorporation methods, among others. For instance, irrigation has been associated with increased SOC in California and other arid/semi-arid croplands, especially in upper surface soil (Mitchell et al. 2017; Emde et al. 2021; Ball et al. 2023). While we observed MAOM accrual over time in both control and crushed rock plots in our irrigated sites, it is possible that in other contexts (such as in non-irrigated croplands in other biomes, associated with SOC losses over time) ERW could cause or exacerbate SOC losses. Furthermore, future ERW field trials should not only include comprehensive measurements of different SOM pools with robust baseline data taken before treatment, but also assess these SOM pools over multiple years and across various soil depths, since effects on SOM can vary substantially over time and at different soil depths. For instance, a 19-year field trial in California showed that SOC gains at 0-30 cm from winter cover crops under conventional management were offset by losses at 30-200 cm, resulting in a net carbon loss (Tautges et al. 2019). And, while liming can have negative effects on SOC over shorter time scales (i.e., months to a few years), these effects can change direction and eventually become positive over longer durations (Paradelo et al. 2015). It will be crucial to not only evaluate the longer-term effects of ERW on total SOM, but also to untangle how crushed rock may differentially affect MAOM and POM pools. For example, while MAOM pools may continue to exhibit decreased rates of accrual over time, they may instead ultimately increase in response to higher weathering rates, as is known to occur over the time scale of soil formation (Slessarev et al. 2021), and has also been observed in some shorter-scale microcosm studies (Buss et al. 2024).

Field and laboratory studies are needed to mechanistically address the interactions between crushed rock, OM, and soil microorganisms, to understand the drivers of C gains or losses, and determine which microbial taxa enhance dissolution of crushed rock or formation of MAOM (Buss et al. 2024). Tools like quantitative stable isotope probing (qSIP) can yield more precise insights into which microbial taxa are active in the presence of crushed rock; measurements of community-level traits can help inform which traits may enhance inorganic and organic C drawdown, such as production of extracellular polymeric substances or altered carbon-use efficiency (Hungate et al. 2015; Lybrand et al. 2019; Sokol et al. 2022a). Beyond microbial priming, various biotic and abiotic mechanisms may influence SOM cycling. For example, elevated root-induced priming of OM under ERW could have also contributed to lower MAOM accrual (Fang et al. 2023) if nutrients released from the weathered rock stimulated more root growth and/or more exudation of particular compounds like organic acids (Keiluweit et al. 2015). Co-applying organic amendments with crushed rock may offer potential benefits in mitigating or reversing negative impacts on soil organic matter (SOM) pools. Identifying the most suitable blend of crushed rock and OM additions may hold promise for optimizing microbial-mediated inorganic and organic C removal (Corbett et al. 2024).

Overall, our results highlight that it will be critical for future studies to monitor SOM stocks, especially over long enough periods to realistically assess net C removal and effects of ERW on soil fertility. As more data becomes available, the effects of ERW on organic matter pools should be included in C accounting frameworks and models to calculate net C removal. Coupling measurements of both inorganic and organic C data – especially in field trials that span diverse soil types, crop types, and climate regions – is crucial for parameterizing the next generation of models that integrate inorganic C dynamics via both rock weathering and OM cycling. Moreover, this data will be vital to inform measurement, reporting, and verification standards. Amidst growing investment in ERW as a scalable C removal approach for drawdown of excess atmospheric CO_2_, the full suite of effects on the soil system must be evaluated before ERW is deployed at broad scales.

## Supporting information

Supplementary Informmation

## ACKNOWLEDGEMENTS

We thank Jessica Wollard at Lawrence Livermore National Laboratory (LLNL) for assistance with DNA extractions and sequencing, Rachel Lindvall at LLNL for ICP-MS analysis, and Brad Erkkila at Yale University for elemental and isotopic analyses. We also thank the staff at Bowles Farming Company (Merced-1 and Merced-2 sites), as well as the University of California, Davis Campbell Tract staff at the Yolo site, who helped extensively with the establishment and maintenance of field trials. Specialty Granules Inc. provided the meta-basalt crushed rock for the field trials. Work at LLNL was supported by the LLNL LDRD Program (22-LW-022; 24-SI-002), and was performed under the auspices of the DOE, Contract DE-AC52-07NA27344. Research at all field sites was supported by a California Strategic Growth Council grant CCR20007.

## REFERENCES

Apprill A, McNally S, Parsons R, Weber L (2015) Minor revision to V4 region SSU rRNA 806R gene primer greatly increases detection of SAR11 bacterioplankton. Aquat Microb Ecol 75:129–137. 10.3354/ame01753

Ball KR, Malik AA, Muscarella C, Blankinship JC (2023) Irrigation alters biogeochemical processes to increase both inorganic and organic carbon in arid-calcic cropland soils. Soil Biology and Biochemistry 187:109189. 10.1016/j.soilbio.2023.109189

Bates D, Mächler M, Bolker B, Walker S (2015) Fitting linear mixed-effects models using lme4. Journal of Statistical Software 67:1–48. 10.18637/jss.v067.i01

Beerling DJ, Kantzas EP, Lomas MR, et al (2020) Potential for large-scale CO2 removal via enhanced rock weathering with croplands. Nature 583:242–248. 10.1038/s41586-020-2448-9

Beerling DJ, Leake JR, Long SP, et al (2018) Farming with crops and rocks to address global climate, food and soil security. Nature Plants 4:138–147. 10.1038/s41477-018-0108-y

Bradford MA, Fierer N, Reynolds JF (2008) Soil carbon stocks in experimental mesocosms are dependent on the rate of labile carbon, nitrogen and phosphorus inputs to soils. Functional Ecology 22:964–974. 10.1111/j.1365-2435.2008.01404.x

Buss W, Hasemer H, Ferguson S, Borevitz J (2024) Stabilisation of soil organic matter with rock dust partially counteracted by plants. Global Change Biology 30:e17052. 10.1111/gcb.17052

Callahan BJ, McMurdie PJ, Rosen MJ, et al (2016) DADA2: High-resolution sample inference from Illumina amplicon data. Nat Methods 13:581–583. 10.1038/nmeth.3869

Cambardella CA, Elliott ET (1992) Particulate soil organic matter changes across a grassland cultivation sequence. Soil Science Society of America Journal 56:777–783. 10.2136/sssaj1992.03615995005600030017x

Chan KY, Heenan DP (1999) Lime-induced loss of soil organic carbon and effect on aggregate stability. Soil Science Society of America Journal 63:1841–1844. 10.2136/sssaj1999.6361841x

Corbett TDW, Westholm M, Rosling A, et al (2024) Organic carbon source controlled microbial olivine dissolution in small-scale flow-through bioreactors, for CO2 removal. npj Mater Degrad 8:1–13. 10.1038/s41529-024-00454-w

Dietzen C, Harrison R, Michelsen-Correa S (2018) Effectiveness of enhanced mineral weathering as a carbon sequestration tool and alternative to agricultural lime: An incubation experiment. International Journal of Greenhouse Gas Control 74:251–258. 10.1016/j.ijggc.2018.05.007

Emde D, Hannam KD, Most I, et al (2021) Soil organic carbon in irrigated agricultural systems: A meta-analysis. Global Change Biology 27:3898–3910. 10.1111/gcb.15680

Fang Q, Lu A, Hong H, et al (2023) Mineral weathering is linked to microbial priming in the critical zone. Nat Commun 14:345. 10.1038/s41467-022-35671-x

Feibelman T, Bayman P, Cibula WG (1994) Length variation in the internal transcribed spacer of ribosomal DNA in chanterelles. Mycological Research 98:614–618. 10.1016/S0953-7562(09)80407-3

Fierer N, Schimel JP, Holden PA (2003) Variations in microbial community composition through two soil depth profiles. Soil Biology and Biochemistry 35:167–176. 10.1016/S0038-0717(02)00251-1

Fuss S, Lamb WF, Callaghan MW, et al (2018) Negative emissions—Part 2: Costs, potentials and side effects. Environ Res Lett 13:063002. 10.1088/1748-9326/aabf9f

Gillman GP (1980) The Effect of Crushed Basalt Scoria on the Cation Exchange Properties of a Highly Weathered Soil. Soil Science Society of America Journal 44:465–468. 10.2136/sssaj1980.03615995004400030005x

Goll DS, Ciais P, Amann T, et al (2021) Potential CO2 removal from enhanced weathering by ecosystem responses to powdered rock. Nat Geosci 14:545–549. 10.1038/s41561-021-00798-x

Guo F, Wang Y, Zhu H, et al (2023) Crop productivity and soil inorganic carbon change mediated by enhanced rock weathering in farmland: A comparative field analysis of multi-agroclimatic regions in central China. Agricultural Systems 210:103691. 10.1016/j.agsy.2023.103691

Gutierrez C, Holland P, O’Neal M (2015) Preliminary Geologic Map of the Ione 7.5’ Quadrangle, Amador County, CA, Version 1.0. California Department of Conservation California Geological Survey. https://www.conservation.ca.gov/cgs/rgm/preliminary

Haque F, Santos RM, Chiang YW (2020a) Optimizing Inorganic Carbon Sequestration and Crop Yield With Wollastonite Soil Amendment in a Microplot Study. Frontiers in Plant Science 11:1012. 10.3389/fpls.2020.01012

Haque F, Santos RM, Chiang YW (2020b) CO2 sequestration by wollastonite-amended agricultural soils – An Ontario field study. International Journal of Greenhouse Gas Control 97:103017. 10.1016/j.ijggc.2020.103017

Haque F, Santos RM, Dutta A, et al (2019) Co-Benefits of Wollastonite Weathering in Agriculture: CO2 Sequestration and Promoted Plant Growth. ACS Omega 4:1425–1433. 10.1021/acsomega.8b02477

Heckman K, Hicks Pries CE, Lawrence CR, et al (2022) Beyond bulk: Density fractions explain heterogeneity in global soil carbon abundance and persistence. Global Change Biology 28:1178–1196. 10.1111/gcb.16023

Holland P (2016) Preliminary Geologic Map of the Irish Hill 7.5’ Quadrangle, Amador County, CA, Version 1.0. California Department of Conservation California Geological Survey. https://www.conservation.ca.gov/cgs/rgm/preliminary

Holzer I, Sokol NW, Slessarev E, et al (2023a) Quantifying enhanced weathering. https://carbonplan.org/research/ew-quantification-explainer. Accessed 15 Nov 2023

Holzer IO, Nocco MA, Houlton BZ (2023b) Direct evidence for atmospheric carbon dioxide removal via enhanced weathering in cropland soil. Environ Res Commun 5:101004. 10.1088/2515-7620/acfd89

Huang Y, Song X, Wang Y-P, et al (2024) Size, distribution, and vulnerability of the global soil inorganic carbon. Science 384:233–239. 10.1126/science.adi7918

Hungate BA, Mau RL, Schwartz E, et al (2015) Quantitative Microbial Ecology through Stable Isotope Probing. Appl Environ Microbiol 81:7570–7581. 10.1128/AEM.02280-15

Kallenbach CM, Grandy AS, Frey SD, Diefendorf AF (2015) Microbial physiology and necromass regulate agricultural soil carbon accumulation. Soil Biology and Biochemistry 91:279–290. 10.1016/j.soilbio.2015.09.005

Kantola IB, Blanc-Betes E, Masters MD, et al (2023) Improved net carbon budgets in the US Midwest through direct measured impacts of enhanced weathering. Global Change Biology 29:7012–7028. 10.1111/gcb.16903

Kantola IB, Masters MD, Beerling DJ, et al (2017) Potential of global croplands and bioenergy crops for climate change mitigation through deployment for enhanced weathering. Biology Letters 13:20160714. 10.1098/rsbl.2016.0714

Keiluweit M, Bougoure JJ, Nico PS, et al (2015) Mineral protection of soil carbon counteracted by root exudates. Nature Clim Change 5:588–595. 10.1038/nclimate2580

Kelland ME, Wade PW, Lewis AL, et al (2020) Increased yield and CO2 sequestration potential with the C4 cereal Sorghum bicolor cultivated in basaltic rock dust-amended agricultural soil. Global Change Biology 26:3658–3676. 10.1111/gcb.15089

Köhler P, Hartmann J, Wolf-Gladrow DA (2010) Geoengineering potential of artificially enhanced silicate weathering of olivine. PNAS 107:20228–20233. 10.1073/pnas.1000545107

Komada T, Anderson MR, Dorfmeier CL (2008) Carbonate removal from coastal sediments for the determination of organic carbon and its isotopic signatures, δ13C and Δ14C: comparison of fumigation and direct acidification by hydrochloric acid. Limnology and Oceanography: Methods 6:254–262. 10.4319/lom.2008.6.254

Kowalenko CG, Ihnat M (2013) Residual effects of combinations of limestone, zinc and manganese applications on soil and plant nutrients under mild and wet climatic conditions. Can J Soil Sci 93:113–125. 10.4141/cjss2011-044

Liang C, Schimel JP, Jastrow JD (2017) The importance of anabolism in microbial control over soil carbon storage. Nature Microbiology 2:1–6. 10.1038/nmicrobiol.2017.105

Liang Y, Rillig MC, Chen HYH, et al (2024) Soil pH drives the relationship between the vertical distribution of soil microbial biomass and soil organic carbon across terrestrial ecosystems: A global synthesis. CATENA 238:107873. 10.1016/j.catena.2024.107873

Love MI, Huber W, Anders S (2014) Moderated estimation of fold change and dispersion for RNA-seq data with DESeq2. Genome Biology 15:550. 10.1186/s13059-014-0550-8

Lybrand RA, Austin JC, Fedenko J, et al (2019) A coupled microscopy approach to assess the nano-landscape of weathering. Scientific Reports 9:5377. 10.1038/s41598-019-41357-0

Malik AA, Puissant J, Buckeridge KM, et al (2018) Land use driven change in soil pH affects microbial carbon cycling processes. Nat Commun 9:3591. 10.1038/s41467-018-05980-1

Martin M (2011) Cutadapt removes adapter sequences from high-throughput sequencing reads. EMBnet.journal 17:10–12. 10.14806/ej.17.1.200

McMurdie PJ, Holmes S (2013) phyloseq: An R package for reproducible interactive analysis and graphics of microbiome census data. PLOS ONE 8:e61217. 10.1371/journal.pone.0061217

Min K, Slessarev E, Kan M, et al (2021) Active microbial biomass decreases, but microbial growth potential remains similar across soil depth profiles under deeply-vs. shallow-rooted plants. Soil Biology and Biochemistry 162:108401. 10.1016/j.soilbio.2021.108401

Minx JC, Lamb WF, Callaghan MW, et al (2018) Negative emissions—Part 1: Research landscape and synthesis. Environ Res Lett 13:063001. 10.1088/1748-9326/aabf9b

Mitchell JP, Shrestha A, Mathesius K, et al (2017) Cover cropping and no-tillage improve soil health in an arid irrigated cropping system in California’s San Joaquin Valley, USA. Soil and Tillage Research 165:325–335. 10.1016/j.still.2016.09.001

Moosdorf N, Renforth P, Hartmann J (2014) Carbon dioxide efficiency of ferrestrial enhanced weathering. Environ Sci Technol 48:4809–4816. 10.1021/es4052022

Nilsson RH, Larsson K-H, Taylor AFS, et al (2019) The UNITE database for molecular identification of fungi: handling dark taxa and parallel taxonomic classifications. Nucleic Acids Research 47:D259–D264. 10.1093/nar/gky1022

Parada AE, Needham DM, Fuhrman JA (2016) Every base matters: assessing small subunit rRNA primers for marine microbiomes with mock communities, time series and global field samples. Environmental Microbiology 18:1403–1414. 10.1111/1462-2920.13023

Paradelo R, Virto I, Chenu C (2015) Net effect of liming on soil organic carbon stocks: A review. Agriculture, Ecosystems & Environment 202:98–107. 10.1016/j.agee.2015.01.005

Quast C, Pruesse E, Yilmaz P, et al (2013) The SILVA ribosomal RNA gene database project: improved data processing and web-based tools. Nucleic Acids Research 41:D590–D596. 10.1093/nar/gks1219

Renforth P (2012) The potential of enhanced weathering in the UK. International Journal of Greenhouse Gas Control 10:229–243. 10.1016/j.ijggc.2012.06.011

Rowley MC, Grand S, Verrecchia ÉP (2018) Calcium-mediated stabilisation of soil organic carbon. Biogeochemistry 137:27–49. 10.1007/s10533-017-0410-1

Scharlemann JP, Tanner EV, Hiederer R, Kapos V (2014) Global soil carbon: understanding and managing the largest terrestrial carbon pool. Carbon Management 5:81–91. 10.4155/cmt.13.77

Schoch CL, Robbertse B, Robert V, et al (2014) Finding needles in haystacks: linking scientific names, reference specimens and molecular data for Fungi. Database 2014:bau061. 10.1093/database/bau061

Shabtai IA, Wilhelm RC, Schweizer SA, et al (2023) Calcium promotes persistent soil organic matter by altering microbial transformation of plant litter. Nat Commun 14:6609. 10.1038/s41467-023-42291-6

Shukla PR, Skea J, Slade R, et al (2022) IPCC, 2022: Climate Change 2022: Mitigation of climate change. Contribution of working group III to the sixth assessment report of the Intergovernmental Panel on Climate Change. Cambridge University Press, Cambridge, UK and New York, NY, USA

Slessarev EW, Chadwick OA, Sokol NW, et al (2021) Rock weathering controls the potential for soil carbon storage at a continental scale. Biogeochemistry. 10.1007/s10533-021-00859-8

Slessarev EW, Lin Y, Jiménez BY, et al (2020) Cellular and extracellular C contributions to respiration after wetting dry soil. Biogeochemistry 2020 147:3 147:307–324. 10.1007/S10533-020-00645-Y

Sokol NW, Foley MM, Blazewicz SJ, et al (2024) The path from root input to mineral-associated soil carbon is dictated by habitat-specific microbial traits and soil moisture. Soil Biology and Biochemistry 193:109367. 10.1016/j.soilbio.2024.109367

Sokol NW, Kuebbing SE, Bradford MA (2017) Impacts of an invasive plant are fundamentally altered by a co-occurring forest disturbance. Ecology 98:2133–2144. 10.1002/ecy.1906

Sokol NW, Slessarev E, Marschmann GL, et al (2022a) Life and death in the soil microbiome: how ecological processes influence biogeochemistry. Nat Rev Microbiol 1–16. 10.1038/s41579-022-00695-z

Sokol NW, Whalen ED, Jilling A, et al (2022b) Global distribution, formation and fate of mineral-associated soil organic matter under a changing climate: A trait-based perspective. Functional Ecology. 10.1111/1365-2435.14040

Strickland MS, Thomason WE, Avera B, et al (2019) Short-term effects of cover crops on soil microbial characteristics and biogeochemical processes across actively managed farms. Agrosystems, Geosciences & Environment 2:180064. 10.2134/age2018.12.0064

Suhrhoff TJ, Reershemius T, Wang J, et al (2024) A tool for assessing the sensitivity of soil-based approaches for quantifying enhanced weathering: a US case study. Front Clim 6:. 10.3389/fclim.2024.1346117

Tautges NE, Chiartas JL, Gaudin ACM, et al (2019) Deep soil inventories reveal that impacts of cover crops and compost on soil carbon sequestration differ in surface and subsurface soils. Global Change Biology 25:3753–3766. 10.1111/gcb.14762

Taylor DL, Walters WA, Lennon NJ, et al (2016) Accurate Estimation of Fungal Diversity and Abundance through Improved Lineage-Specific Primers Optimized for Illumina Amplicon Sequencing. Applied and Environmental Microbiology 82:7217–7226. 10.1128/AEM.02576-16

Uroz S, Calvaruso C, Turpault M-P, Frey-Klett P (2009) Mineral weathering by bacteria: ecology, actors and mechanisms. Trends in Microbiology 17:378–387. 10.1016/j.tim.2009.05.004

Vicca S, Goll DS, Hagens M, et al (2022) Is the climate change mitigation effect of enhanced silicate weathering governed by biological processes? Global Change Biology 28:711– 726. 10.1111/gcb.15993

Yan Y, Dong X, Li R, et al (2023) Wollastonite addition stimulates soil organic carbon mineralization: Evidences from 12 land-use types in subtropical China. CATENA 225:107031. 10.1016/j.catena.2023.107031

