## Supplementary Informmation for "Reduced accrual of mineral-associated organic matter after two years of enhanced rock weathering in cropland soils, though no net losses of soil organic carbon"

**Description**: *This supplementary file contains a description of the experimental design of the three field sites and the site characteristics, the output of all linear mixed effect models, the means and standard errors of SOM pools (including stocks in fall 2021 and two-year changes through time between fall 2019 and fall 2021), as well microbial community analysis output of bacterial and fungal sequence data in fall 2021.*
*(Figures S1, Tables S1- S12.)*

**Figure S1. Experimental design of three enhanced rock weathering field trials in California, USA.** The three sites include the: (a) Yolo site (n = 3), (b) Merced-1 site (n = 5), and (c) Merced-2 site (n = 5). The Yolo site included several other treatments (compost, biochar, rock + compost, rock + biochar, compost + biochar, rock + compost + biochar); this study only included only the rock and control plots. At Yolo, rock plots were divided in half, to include a meta-basalt amendment and an olivine amendment; this study only used the meta-basalt treatment). Further site details are shown in Table S1.

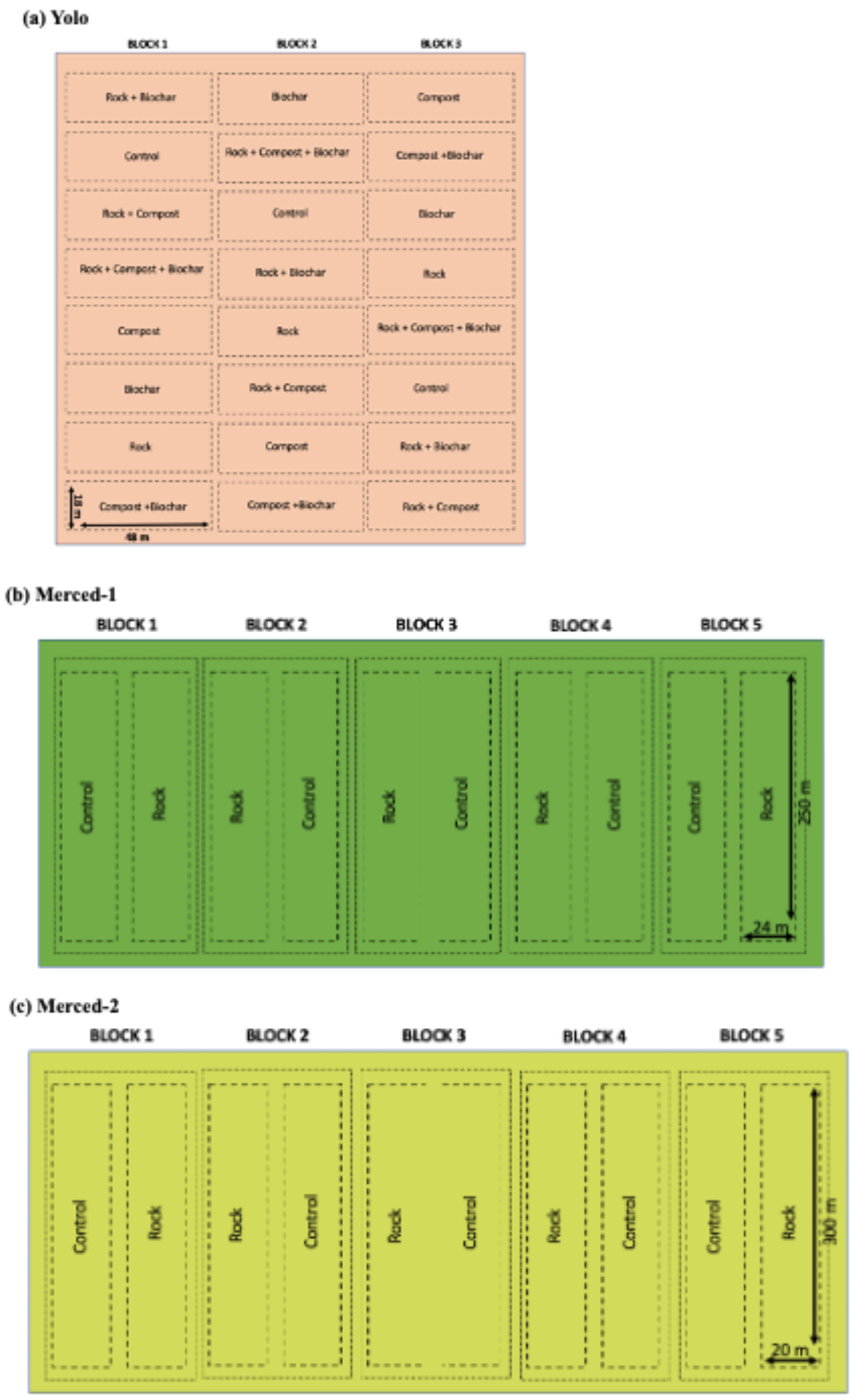

**Table S1. Site details for three enhanced rock weathering field trials at the Yolo, Merced-1, and Merced 2 sites.**

| Site | Yolo | Merced-1 | Merced-2 |
|---|---|---|---|
| **Location** | 38° 31'53.96"N, 121°46'54.15"W | 37° 7'10.099"N, 120°44'38.41"W | |
| **Mean annual precipitation (mm)** | 449 | 174 | |
| **Mean annual temperature (°C)** | 15.6 | 16.7 | |
| **Crop** | Corn | Alfalfa | *Rotation*: Corn (2019), Tomato (2020), Cilantro (2021) |
| **Management** | Conventional | Organic, Conventional | |
| **Irrigation** | Drip (subsurface) | Furrow/Flood | |
| **Site size (ha)** | 5.04 | 6.1 | |
| **Plot size (ha)** | 0.09 (0.045 for plots with meta-basalt rock plots) | 0.6 | |
| **Reps/treatment** | 3 | 5 | 5 |
| **Soil type** | Coarse-loamy, mixed, superactive, nonacid, thermic Mollic Xerofluvents | Fine-loamy, mixed (calcareous), thermic Typic Haplaquolls | Fine-loamy, mixed, montmorillonitic (calcareous), superactive, thermic Typic Epiaqualfs |
| **pH** | 6.7-7.5 | 7.9-8.2 | 7.9-8.2 |
| **USDA soil texture class** | Silt Loam | Clay Loam* | |
| **Soil texture percentages (%)** | 4.0 sand; 61.0 silt; 25.1 clay | 35.4 Sand, 33.6 Silt, 31 Clay* | |
| **Meta-basalt crushed rock application** | 40 t $ha^{-1}$ (Fall 2019 and 2020) | 40 t $ha^{-1}$ (Fall 2019 only) | 40 t $ha^{-1}$ (Fall 2019 and 2020) |
| **Parent Material** | Alluvium from mixed sources | Alluvium from granite | |
| **Soil Series/Order** | Reiff Series/Entisol | Escano Series/Mollisol | Alros Series/Alfisol |

**data is from the Soil Web Series. (Reference: Soil Survey Staff, Natural Resources Conservation Service, United States Department of Agriculture. Web Soil Survey. Available online at http://websoilsurvey.nrcs.usda.gov/ accessed [11/16/2023].)*

**Table S2. Elemental composition of meta-basalt crushed rock applied each year of the study and reported as oxide weight-percent.** The high silica content of the dust results from felsic facies included in the mined and crushed material (Holzer et al, 2023b). Method detection limits are reported parenthetically after each oxide. LOI = loss on ignition.

| **Oxide (%)** | **Yolo, 2019** | **Merced-1 and -2, 2019** | **Yolo, 2020** | **Merced-2, 2020** | **Yolo, 2021** |
|---|---|---|---|---|---|
| **$SiO_2$ (0.01)** | 65.92 | 65.63 | 61.66 | 65.51 | 66.78 |
| **$Al_2O_3$ (0.01)** | 15.22 | 15.42 | 15.05 | 15.61 | 15.11 |
| **$Fe_2O_3$ (0.04)** | 4.9 | 4.86 | 6.52 | 4.85 | 4.43 |
| **MgO (0.01)** | 1.66 | 1.7 | 3.27 | 1.84 | 1.52 |
| **CaO (0.01)** | 3.74 | 3.79 | 5.19 | 3.9 | 3.85 |
| **$Na_2O$ (0.01)** | 4.21 | 4.2 | 3.36 | 3.56 | 3.85 |
| **$K_2O$ (0.01)** | 1.46 | 1.43 | 1.28 | 1.54 | 1.5 |
| **$TiO_2$ (0.01)** | 0.48 | 0.48 | 0.54 | 0.52 | 0.47 |
| **$P_2O_5$ (0.01)** | 0.17 | 0.14 | 0.15 | 0.17 | 0.15 |
| **MnO (0.01)** | 0.14 | 0.13 | 0.16 | 0.12 | 0.09 |
| **$Cr_2O_3$ (0.002)** | 0.004 | 0.008 | 0.014 | 0.005 | 0.01 |
| **BaO (0.01)** | 0.05 | 0.06 | 0.06 | 0.06 | 0.05 |
| **LOI (0.1)** | 1.9 | 2 | 2.6 | 2.2 | 2.1 |
| **Sum** | 99.9 | 99.89 | 99.91 | 99.91 | 99.94 |

**Table S3. Pre-treatment baseline concentrations of upper surface soil SOM pools (at 0-10 cm) measured in fall 2019 before the experimental period.** Plot-specific baseline measurements were taken at two of the three sites (Yolo and Merced-2) before treatments were applied ( 'pre-trt. Ctr.' = pre-treatment control plots; 'Pre-trt. Rock'= pre-treatment crushed rock plots). Shown below are means ± standard error (SE) of SOM pool concentrations, including total soil organic C (SOC), total soil organic N (SON), mineral-associated organic matter (MAOM) C and N, and particulate organic matter (POM) C and N. Percent difference (% diff.) is the average difference in the plot-specific concentration of an SOM pool in pre-treatment rock and control plots. At Yolo, baseline data indicated there were significant initial differences between treatment plots for several SOM pools ($p < 0.05$; indicated with an *), which were accounted as a correction factor when measuring % differences in fall 2021 SOM stocks (Table S5). At Merced-2, there was no significant difference between baseline values of treatment plots ($p > 0.1$). Negative % difference values indicate that SOM concentrations are lower in the experimental plots where rock would be applied versus control plots. *N* = 5 at Merced-2, *n* =3 at Yolo.

| Site | Trt. | SOC ± SE (mg C g soil$^{-1}$) | % Diff. ± SE | SON ± SE (mg C g soil$^{-1}$) | % Diff. ± SE | MAOM-C ± SE (mg C g soil$^{-1}$) | % Diff. ± SE | MAOM-N ± SE (mg C g soil$^{-1}$) | % Diff. ± SE | POM-C ± SE (mg C g soil$^{-1}$) | % Diff. ± SE | POM-N ± SE (mg C g soil$^{-1}$) | % Diff. ± SE |
|---|---|---|---|---|---|---|---|---|---|---|---|---|---|
| Merced-2 | Pre-trt. Ctr. | 9.04 ± 0.3 | NS | 1.07 ± 0.04 | NS | 6.59 ± 0.16 | NS | 0.85 ± 0.03 | NS | 2.45 ± 0.18 | NS | 0.21 ± 0.02 | NS |
| | Pre-trt. Rock | 8.84 ± 0.41 | | 1.02 ± 0.03 | | 6.39 ± 0.19 | | 0.83 ± 0.02 | | 2.45 ± 0.26 | | 0.19 ± 0.02 | |
| Yolo | Pre-trt. Ctr. | 6.57 ± 0.32 | -14 ± 5* | 0.76 ± 0.03 | -11 ± 4* | 5.4 ± 0.3 | -17 ± 4* | 0.61 ± 0.03 | -13 ± 4* | 1.19 ± 0.1 | -3 ± 10 | 0.15 ± 0.001 | -6 ± 4 |
| | Pre-trt. Rock | 5.66 ± 0.21 | | 0.68 ± 0.02 | | 4.5 ± 0.2 | | 0.53 ± 0.02 | | 1.16 ± 0.04 | | 0.14 ± 0.001 | |

**Table S4. Modeled regression coefficients of the effect of site and crushed rock on soil organic matter and inorganic carbon stocks in three cropland enhanced rock weathering field trials.** Shown below are coefficient estimates (± standard error, SE) from linear mixed effect models, which included the effect of crushed rock and site on SOM stocks in Fall 2021. *N = 13* per treatment across all field trials. SOC = total soil organic C; SON = total soil organic N; MAOM = mineral-associated organic matter; POM = particulate organic matter, SIC = soil inorganic C. *(p < 0.1 is considered marginally significant and indicated by a single asterisks *; p < 0.05 is considered significant and indicated by two asterisks **).*

| | **Coefficient estimate ± SE** | **p value** |
|---|---|---|
| ***(a)*** *Total SOC (0-10 cm)* | | |
| Intercept | 1339 ± 65 | <0.0001** |
| Site (Merced-2) | 386 ± 61 | <0.0001** |
| Site (Yolo) | -370 ± 73 | <0.0004** |
| Treatment (crushed rock) | -140 ± 53 | 0.02** |
| ***(b)*** *Total SOC (10-30 cm)* | | |
| Intercept | 1730 ± 71 | <0.0001** |
| Site (Merced-2) | 1000 ± 86 | <0.0001** |
| Site (Yolo) | -149 ± 99 | 0.15 |
| Treatment (crushed rock) | -99 ± 75 | 0.2 |
| ***(c)*** *Total SON (0-10 cm)* | | |
| Intercept | 151 ± 8 | <0.0001** |
| Site (Merced-2) | 45 ± 7 | <0.0001** |
| Site (Yolo) | -49 ± 9 | 0.0002** |
| Treatment (crushed rock) | -19 ± 6 | 0.007** |
| ***(d)*** *Total SON (10-30 cm)* | | |
| Intercept | 188 ± 9 | <0.0001** |
| Site (Merced-2) | 120 ± 10 | <0.0001** |
| Site (Yolo) | 0.7 ± 12 | 0.96 |
| Treatment (crushed rock) | -3 ± 9 | 0.74 |
| ***(e)*** *MAOM-C (0-10 cm)* | | |
| Intercept | 952 ± 46 | <0.0001** |
| Site (Merced-2) | 251 ± 42 | <0.0001** |
| Site (Yolo) | -263 ± 51 | 0.0004** |
| Treatment (crushed rock) | -159 ± 37 | 0.0001** |
| **(f)** *MAOM-C (10-30 cm)* | | |
| Intercept | 1395 ± 61 | <0.0001** |
| Site (Merced-2) | 660 ± 74 | <0.0001** |
| Site (Yolo) | -65 ± 85 | 0.45 |
| Treatment (crushed rock) | -68 ± 65 | 0.30 |
| **(g)** *MAOM-N (0-10 cm)* | | |
| Intercept | 116 ± 6 | <0.0001** |
| Site (Merced-2) | 34 ± 5 | <0.0001** |
| Site (Yolo) | -39 ± 7 | <0.0001** |
| Treatment (crushed rock) | -17 ± 5 | 0.001** |

| | | |
|---|---|---|
| **(h)** *MAOM-N (10-30 cm)* | | |
| Intercept | 165 ± 8 | <0.0001** |
| Site (Merced-2) | 83 ± 10 | <0.0001** |
| Site (Yolo) | -12 ± 11 | 0.27 |
| Treatment (crushed rock) | -7 ± 8 | 0.41 |
| **(i)** *POM-C (0-10 cm)* | | |
| Intercept | 387 ± 32 | <0.0001** |
| Site (Merced-2) | 135 ± 36 | 0.0015** |
| Site (Yolo) | -107 ± 42 | 0.02** |
| Treatment (crushed rock) | 19 ± 31 | 0.6 |
| **(j)** *POM-C (10-30 cm)* | | |
| Intercept | 335 ± 39 | <0.0001** |
| Site (Merced-2) | 341 ± 46 | <0.0001** |
| Site (Yolo) | -78 ± 53 | 0.16 |
| Treatment (crushed rock) | -31 ± 40 | 0.45 |
| **(k)** *POM-N (0-10 cm)* | | |
| Intercept | 34 ± 3 | <0.0001** |
| Site (Merced-2) | 11 ± 3 | <0.0001** |
| Site (Yolo) | -8 ± 3 | 0.02** |
| Treatment (crushed rock) | -2 ± 2 | 0.52 |
| **(l)** *POM-N (10-30 cm)* | | |
| Intercept | 23 ± 2 | <0.0001** |
| Site (Merced-2) | 37 ± 3 | <0.0001** |
| Site (Yolo) | 13 ± 3 | <0.0001** |
| Treatment (crushed rock)) | 4 ± 2 | 0.12 |
| **(m)** *DOC (0-10 cm)* | | |
| Intercept | 14 ± 2 | <0.0001** |
| Site (Merced-2) | 14 ± 2 | <0.0001** |
| Site (Yolo) | 13 ± 2 | <0.0001** |
| Treatment (crushed rock) | 0.07 ± 1 | 0.96 |
| **(n)** *DOC (10-30 cm)* | | |
| Intercept | 11 ± 1 | <0.0001** |
| Site (Merced-2) | 6 ± 1 | <0.0001** |
| Site (Yolo) | -0.01 ± 1 | 0.992 |
| Treatment (crushed rock) | -0.002 ± 1 | 0.998 |
| **(o)** *SIC (0-10 cm)* | | |
| Intercept | 59 ± 34 | 0.096 |
| Site (Merced-2) | 124 ± 39 | 0.006** |
| Site (Yolo) | -22 ± 45 | 0.63 |
| Treatment (crushed rock) | 13 ± 34 | 0.70 |
| **(p)** *SIC (10-30 cm)* | | |
| Intercept | 270 ± 64 | 0.001** |
| Site (Merced-2) | -5.1 ± 66 | 0.94 |
| Site (Yolo) | -218 ± 78 | 0.011** |
| Treatment (crushed rock) | 89 ± 57 | 0.14 |

**Table S5. Mean stock sizes of SOM pools in upper surface soil (0-10 cm) of plots with crushed rock and unamended control plots in fall 2021 at three enhanced rock weathering field trials**. Absolute difference ('Diff') is the difference between the mean of the crushed rock plots minus the mean of the control plots at a given site. Percent difference (% diff.) is calculated as the difference between rock and control plots, divided by the value of the control plots at a given site. At Yolo, pre-treatment baseline data (from 2019, before crushed rock application) indicated there were significant initial differences between treatment plots ($p < 0.05$; see Table S3), though there was no significant difference in baseline concentrations at Merced-2. At Yolo, a correction factor was thus applied to account for baseline differences when estimating the % difference between stock sizes in control versus rock plots (see Methods). Standard errors for absolute differences and percent differences are calculated using gaussian error propagation. Negative % difference values indicate that SOM stocks are lower in rock versus control plots.

| **Site** | **Trt.** | **SOC Stock ± SE** (g C $m^{-2}$ to 10 cm) | *Diff. ± SE (g C $m^{-2}$ to 10 cm)* | *% Diff ± SE* | **SON Stock ± SD (g N $m^{-2}$ to 10 cm)** | *Diff. (g N $m^{-2}$ to 10 cm)* | *% Diff. ± SE* | **MAOM C Stock ± SD (g C $m^{-2}$ to 10 cm)** | *Diff ± SE (g C $m^{-2}$ to 10 cm).* | *% Diff ± SE* | **MAOM N Stock ± SD (g N $m^{-2}$ to 10 cm)** | *Diff. ± SE. (g N $m^{-2}$ to 10 cm)* | *% Diff ± SE* | **POM C Stock ± SD (g C $m^{-2}$ to 10 cm)** | *Diff. ± SE. (g C $m^{-2}$ to 10 cm)* | *% Diff ± SE* | **POM N Stock ± SD (g N $m^{-2}$ to 10 cm)** | *Diff. ± SE. (g N $m^{-2}$ to 10 cm)* | *% Diff. ± SE* |
|---|---|---|---|---|---|---|---|---|---|---|---|---|---|---|---|---|---|---|---|
| **Merced-1** | Ctr | 1326 ± 44 | -116 ± 84 | -9 ± 4 | 148 ± 5 | -14 ± 8 | -10 ± 4 | 949 ± 49 | -152 ± 61 | -16 ± 5 | 116 ± 5 | -16 ± 7 | -14 ± 4 | 378 ± 19 | 37 ± 49 | 10 ± 8 | 33 ± 1 | -1 ± 4 | -3 ± 7 |
| | Rock | 1211 ± 71 | | | 134 ± 7 | | | 796 ± 36 | | | 100 ± 4 | | | 415 ± 44 | | | 34 ± 3 | | |
| **Merced-2** | Ctr | 1726 ± 102 | -141± 146 | -8 ± 6 | 198 ± 14 | -23 ± 19 | -11 ± 7 | 1189 ± 73 | -132 ± 96 | -11 ± 6 | 151 ± 11 | -17 ± 14 | -11 ± 8 | 536 ± 39 | -10 ± 66 | -2 ± 9 | 47 ± 3 | 6 ± 6 | 13 ± 8 |
| | Rock | 1584 ± 105 | | | 175 ± 13 | | | 1057 ± 63 | | | 134 ± 9 | | | 527 ± 54 | | | 41 ± 5 | | |
| **Yolo** | Ctr | 1001 ± 58 | -179 ± 62 | -18 ± 5 | 108 ± 5 | -22 ± 6 | -20 ± 4 | 730 ± 47 | -217 ± 59 | -30 ± 12 | 82 ± 7 | -22 ± 6 | -27 ± 4 | 271 ± 17 | 38 ± 54 | 14 ± 12 | 27 ± 1 | -1 ± 2 | -4 ± 5 |
| | Rock | 822 ± 20 | | | 86 ± 3 | | | 513 ± 36 | | | 60 ± 7 | | | 309 ± 51 | | | 26 ± 2 | | |
| | *Baseline % Diff.* | | | *-14 ± 5* | | | *-11 ± 4* | | | *-17 ± 4* | | | *-13 ± 4* | | | *-3 ± 10* | | | *-6 ± 4* |
| | Corrected % Diff. | | | -4 ± 6 | | | -10 ± 6 | | | -13 ±12 | | | *-13 ± 6* | | | *17 ± 16* | | | *3 ± 6* |

**Table S6. Modeled regression coefficients of the effect of site and crushed rock on plot-specific two-year change in soil organic matter concentrations in enhanced rock weathering field trials.** Shown below are coefficient estimates (± standard error, SE) from linear mixed effect models, which included the effect of crushed rock and site on the plot-specific change in soil organic matter pools over the 2-year experimental period (fall 2019 – fall 2021). This dataset includes the two sites where baseline data was available (Yolo and Merced-2). SOC = total soil organic C; SON = total soil organic N; MAOM = mineral-associated organic matter; POM = particulate organic matter. *($p < 0.1$ is considered marginally significant and indicated by a single asterisks *; $p < 0.05$ is considered significant and indicated by two asterisks **).*

| | **Coefficient estimate ± SE** | **p value** |
|---|---|---|
| ***(a)*** *2-year change in SOC (0-10 cm)* | | |
| Intercept | 1.2 ± 0.3 | <0.002** |
| Site (Yolo) | 0.2 ± 0.4 | 0.66 |
| Treatment (crushed rock) | -0.3 ± 0.4 | 0.51 |
| ***(b)*** *2-year change in SOC (10-30 cm)* | | |
| Intercept | 2.2 ± 0.4 | <0.004** |
| Site (Yolo) | -1.7 ± 0.3 | 0.0007** |
| Treatment (crushed rock) | -0.07 ± 0.3 | 0.81 |
| ***(c)*** *2-year change in SON (0-10 cm)* | | |
| Intercept | 0.12 ± 0.05 | 0.05* |
| Site (Yolo) | -0.05 ± 0.04 | 0.3 |
| Treatment (crushed rock) | -0.03 ± 0.04 | 0.5 |
| ***(d)*** *2-year change in SON (10-30 cm)* | | |
| Intercept | 0.18 ± 0.03 | 0.0001** |
| Site (Yolo) | -0.22 ± 0.04 | 0.0001** |
| Treatment (crushed rock) | 0.03 ± 0.04 | 0.46 |
| ***(e)*** *2-year change in MAOM-C (0-10 cm)* | | |
| Intercept | 0.68 ± 0.2 | 0.005** |
| Site (Yolo) | -0.59 ± 0.2 | 0.017** |
| Treatment (crushed rock) | -0.67 ± 0.2 | 0.004** |
| Treatment (rock)*Site (Yolo) | 0.55 | 0.07* |
| **(f)** *2-year change in MAOM-C (10-30 cm)* | | |
| Intercept | 1 ± 0.4 | 0.04** |
| Site (Yolo) | -0.7 ± 0.3 | 0.02** |
| Treatment (crushed rock) | 0.2 ± 0.2 | 0.4 |
| **(g)** *2-year change in MAOM-N (0-10 cm)* | | |
| Intercept | 0.06 ± 0.04 | 0.2 |
| Site (Yolo) | -0.04 ± 0.03 | 0.0996* |
| Treatment (crushed rock) | -0.04 ± 0.02 | 0.10 |
| **(h)** *2-year change in MAOM-N (10-30 cm)* | | |
| Intercept | 0.08 ± 0.03 | 0.008** |
| Site (Yolo) | -0.12 ± 0.03 | 0.003** |
| Treatment (crushed rock) | 0.03 ± 0.03 | 0.44 |

| | | |
|---|---|---|
| **(i)** *2-year change in POM-C (0-10 cm)* | | |
| Intercept | 0.7 ± 0.2 | 0.01** |
| Site (Yolo) | 0.5 ± 0.3 | 0.2 |
| Treatment (crushed rock) | 0.2 ± 0.3 | 0.5 |
| **(j)** *2-year change in POM-C (10-30 cm)* | | |
| Intercept | 1.2 ± 0.2 | <0.0001** |
| Site (Yolo) | -1.1 ± 0.2 | 0.0006** |
| Treatment (crushed rock) | -0.3 ± 0.2 | 0.25 |
| **(k)** *2-year change in POM-N (0-10 cm)* | | |
| Intercept | 0.07 ± 0.02 | 0.003** |
| Site (Yolo) | -0.01 ± 0.02 | 0.9 |
| Treatment (crushed rock) | 0.01 ± 0.02 | 0.6 |
| **(l)** *2-year change in POM-N (10-30 cm)* | | |
| Intercept | 0.1 ± 0.01 | <0.0001** |
| Site (Yolo) | -0.1 ± 0.02 | <0.0001** |
| Treatment (crushed rock) | 0.009 ± 0.01 | 0.55 |
| **(m)** *2-year change in soil inorganic C (0-10 cm)* | | |
| Intercept | 0.4 ± 0.3 | 0.28 |
| Site (Yolo) | -0.38 ± 0.26 | 0.17 |
| Treatment (crushed rock) | 0.68 ± 0.23 | 0.017 ** |
| **(n)** *2-year change in soil inorganic C (10-30 cm)* | | |
| Intercept | 0.31 ± 0.22 | 0.18 |
| Site (Yolo) | -0.42 ± 0.28 | 0.16 |
| Treatment (crushed rock) | -0.08 ± 0.27 | 0.76 |

**Table S7. Plot-specific two-year change in SOM pools in the surface soil (0-10 cm) of plots with crushed rock versus control plots.** This dataset includes the two sites where pre-treatment baseline data was available (Yolo and Merced-2): the two-year change is calculated as the plot-specific change in SOM concentrations between Fall 2019 (baseline pre-treatment measurements) and Fall 2021 (see Methods). Absolute difference (‘Diff’) is the difference between the mean of the crushed rock plots minus the mean of the control plots at a given site. Standard errors for absolute differences calculated using gaussian error propagation. There were no significant differences at 10-30 cm depth (Table S5).

| Site | Trt. | 2-year change in total SOC ± SE (mg C g soil$^{-1}$) | Diff. ± SE (mg C g soil$^{-1}$) | 2-year change in total SON ± SE (mg N g soil$^{-1}$) | Diff. ± SE (mg C g soil$^{-1}$) | 2-year change in MAOM-C ± SE (mg C g soil$^{-1}$) | Diff. ± SE (mg C g soil$^{-1}$) | 2-year change in MAOM-N ± SE (mg C g soil$^{-1}$) | Diff. ± SE (mg N g soil$^{-1}$) | 2-year change in POM-C ± SE (mg C g soil$^{-1}$) | Diff. ± SE (mg C g soil$^{-1}$) | 2-year change in POM-N ± SE (mg C g soil$^{-1}$) | Diff. ± SE mg N g soil$^{-1}$) | 2-year change in total soil inorganic C (mg C g soil$^{-1}$) | Diff. ± SE (mg C g soil$^{-1}$) |
|---|---|---|---|---|---|---|---|---|---|---|---|---|---|---|---|
| Merced-2 | Ctr. | 1.51 ± 0.7 | –0.8 ± 0.41 | 0.14 ± 0.12 | –0.06 ± 0.08 | 0.68 ± 0.05 | -0.67 ± 0.26 | 0.07 ± 0.09 | -0.06 ± 0.06 | 0.84 ± 0.17 | -0.13 ± 0.3 | 0.074 ± 0.8 | -0.01 ± 0.03 | 0.22 ± 0.4 | 0.96 ± 0.02 |
| | Rock | 1.08 ± 1.9 | | 0.08 ± 0.13 | | 0.26 ± 0.01 | | 0.009 ± 0.09 | | 0.71 ± 0.2 | | 0.068 ± 0.7 | | 1.18 ± 0.2 | |
| Yolo | Ctr. | 0.98 ± 0.6 | 0.63 ± 0.51 | 0.05 ± 0.05 | 0.03 ± 0.03 | 0.12 ± 0.3 | -0.12 ± 0.17 | 0.00015 ± 0.04 | -0.01 ± 0.02 | 0.86 ± 0.2 | 0.74 ± 0.6 | 0.05 ± 0.9 | 0.04 ± 0.03 | 0.11 ± 0.2 | 0.21 ± 0.16 |
| | Rock | 1.61 ± 0.98 | | 0.08 ± 0.03 | | 0.004 ± 0.03 | | -0.005 ± 0.02 | | 1.6 ± 0.6 | | 0.09 ± 1.6 | | 0.32 ± 0.15 | |

**Table S8.** $\boldsymbol{\delta^{13}C}$ **and** $\boldsymbol{\delta^{15}N}$ **values of soil organic matter pools in three cropland enhanced rock weathering field trials.** Shown below are means (± standard error, SE) of $\delta^{13}C$ and $\delta^{15}N$ values of mineral-associated organic matter (MAOM), particulate organic matter (POM), and dissolved organic carbon (DOC) pools at 0-10 cm and 10-30 cm in plots with crushed rock and control plots. $N = 5$ Merced-1 and Merced-2; $n = 3$ at Yolo.

| Site | Depth (cm) | Treatment | MAOM $\delta^{13}C$ ± SE | MAOM $\delta^{15}N$ ± SE | POM $\delta^{13}C$ ± SE | POM $\delta^{15}N$ ± SE | DOC d $\delta^{13}C$ ± SE |
|---|---|---|---|---|---|---|---|
| **Merced-1** | 0-10 | Control | -23.92 ± 0.97 | 7.24 ± 0.1 | -26.56 ± 0.58 | 4.99 ± 0.17 | -24.93 ± 0.65 |
| | | Rock | -25.40 ± 0.08 | 7.21 ± 0.1 | -26.48 ± 0.62 | 4.87 ± 0.17 | -24.33 ±1.36 |
| | 10-30 | Control | -24.71 ± 0.6 | 6.93 ± 0.1 | -20.59 ± 1.78 | 5.47 ± 0.11 | -25.54 ± 0.2 |
| | | Rock | -24.85 ± 0.3 | 6.84 ± 0.14 | -25.37 ±1.31 | 5.34 ± 0.18 | -23.57 ± 0.7 |
| **Merced-2** | 0-10 | Control | -23.90 ± 0.34 | 7.98 ± 0.11 | -22.18 ± 0.64 | 5.96 ± 0.04 | -23.34 ±0.21 |
| | | Rock | -24.26 ± 0.17 | 7.76 ± 0.1 | -21.94 ±1.08 | 5.79 ± 0.12 | -22.77 ± 0.26 |
| | 10-30 | Control | -24.03 ± 0.73 | 7.17 ± 0.07 | -21.78 ±1.32 | 5.40 ± 0.12 | -23.82 ±0.16 |
| | | Rock | -24.21 ± 0.46 | 7.29 ± 0.05 | -22.93 ± 1.24 | 5.80 ± 0.07 | -23.88 ±0.15 |
| **Yolo** | 0-10 | Control | -25.07 ± 0.13 | 5.55 ± 0.24 | -23.71 ± 0.26 | 2.31 ± 0.21 | -24.24 ± 0.2 |
| | | Rock | -24.65 ± 0.14 | 5.17 ± 0.29 | -23.76 ± 0.6 | 2.04 ± 0.16 | -23.08 ± 0.68 |
| | 10-30 | Control | -25.43 ± 0.02 | 5.11 ± 0.17 | -25.09 ± 0.18 | 2.25 ± 0.18 | -24.5 ± 0.82 |
| | | Rock | -24.93 ± 0.13 | 5.23 ± 0.2 | -23.81 ± 0.26 | 1.82 ± 0.08 | -21.27 ±0.26 |

**Table S9. Modeled regression coefficients of the effect of site and crushed rock on *$\delta^{13}C$* and *$\delta^{15}N$* of soil organic matter pools in three cropland enhanced rock weathering field trials.** Shown below are coefficient estimates (± standard error, SE) from linear mixed effect models, which included the effect of crushed rock and site on C and N stable isotope ratios of soil organic matter pools. *N = 13* per treatment across all field trials. MAOM = mineral-associated organic matter; POM = particulate organic matter; DOC = dissolved organic carbon. *($p < 0.1$ is considered marginally significant and indicated by a single asterisks *; $p < 0.05$ is considered significant and indicated by two asterisks **).*

| | **Coefficient estimate ± SE** | **p value** |
|---|---|---|
| ***(a)*** *$\delta^{13}C$ MAOM-C (0-10 cm)* | | |
| Intercept | -24.3 ± 0.4 | <0.0001** |
| Site (Merced-2) | -0.58 ± 0.5 | 0.24 |
| Site (Yolo) | -0.20 ± 0.6 | 0.72 |
| Treatment (crushed rock) | -0.61 ± 0.5 | 0.16 |
| **(b)** *$\delta^{13}C$ MAOM-C (10-30 cm)* | | |
| Intercept | -24.8 ± 0.4 | <0.0001** |
| Site (Merced-2) | 0.66 ± 0.5 | 0.2 |
| Site (Yolo) | -0.38 ± 0.5 | 0.5 |
| Treatment (crushed rock) | -0.007 ± 0.4 | 0.96 |
| **(c)** *$\delta^{15}N$ MAOM-N (0-10 cm)* | | |
| Intercept | 7.3 ± 0.1 | <0.0001** |
| Site (Merced-2) | 0.6 ± 0.1 | <0.0001** |
| Site (Yolo) | -1.9 ± 0.2 | <0.0001** |
| Treatment (crushed rock) | -0.2 ± 0.1 | 0.1 |
| **(d)** *$\delta^{15}N$ MAOM-N (10-30 cm)* | | |
| Intercept | 6.8 ± 0.1 | <0.0001** |
| Site (Merced-2) | 0.3 ± 0.1 | 0.004 |
| Site (Yolo) | -1.7 ± 0.1 | <0.0001** |
| Treatment (crushed rock) | 0.04 ± 0.1 | 0.67 |
| **(e)** *$\delta^{13}C$ POM-C (0-10 cm)* | | |
| Intercept | 27 ± 0.6 | <0.0001** |
| Site (Merced-2) | 4.5 ± 0.5 | <0.0001** |
| Site (Yolo) | 3.2 ± 0.6 | <0.0001** |
| Treatment (crushed rock) | 0.1 ± 0.5 | 0.81 |
| **(f)** *$\delta^{13}C$ POM-C (10-30 cm)* | | |
| Intercept | -22.0 ± 1 | <0.0001** |
| Site (Merced-2) | 0.6 ± 1 | 0.6 |
| Site (Yolo) | -1.5 ± 2 | 0.4 |
| Treatment (crushed rock) | -2.0 ± 1 | 0.1 |
| **(g)** *$\delta^{15}N$ POM-N (0-10 cm)* | | |
| Intercept | 5.0 ± 0.1 | <0.0001** |
| Site (Merced-2) | 0.9 ± 0.1 | <0.0001** |
| Site (Yolo) | -2.7 ± 0.2 | <0.0001** |
| Treatment (crushed rock) | -0.1 ± 0.1 | 0.1 |

| | | |
|---|---|---|
| **(h)** *$\delta^{15}$N POM-N (10-30 cm)* | | |
| Intercept | 5.4 ± 0.1 | <0.0001** |
| Site (Merced-2) | 0.2 ± 0.1 | 0.18 |
| Site (Yolo) | -3.3 ± 0.2 | <0.0001** |
| Treatment (crushed rock) | 0.007 ± 0.1 | 0.96 |
| **(i)** *$\delta^{13}$C -DOC (0-10 cm)* | | |
| Intercept | -25.0 ± 0.6 | <0.0001** |
| Site (Merced-2) | 1.6 ± 0.6 | 0.02** |
| Site (Yolo) | 0.8 ± 0.7 | 0.2 |
| Treatment (crushed rock) | 0.7 ± 0.5 | 0.2 |

**Table S10. Modeled regression coefficients of the effect of site and crushed rock on edaphic and microbial variables in three enhanced rock weathering field trials.** Shown below are coefficient estimates (± standard error, SE) from linear mixed effect models, which included the effect of crushed rock and site on different soil variables. *N = 13* per treatment across all field trials. *($p < 0.1$ is considered marginally significant and indicated by a single asterisks *; $p < 0.05$ is considered significant and indicated by two asterisks **).*

| | **Coefficient estimate ± SE** | **p value** |
|---|---|---|
| ***(a)*** *Soil pH (10-30 cm)* | | |
| Intercept | 8.1 ± 0.05 | <0.0001** |
| Site (Merced-2) | -0.5 ± 0.05 | <0.0001** |
| Site (Yolo) | -0.6 ± 0.07 | <0.0001** |
| Treatment (crushed rock) | 0.1 ± 0.05 | 0.2 |
| ***(b)*** *Electrical Conductivity (0-10 cm)* | | |
| Intercept | 378 ± 36 | <0.0001** |
| Site (Merced-2) | 266 ± 38 | <0.0001** |
| Site (Yolo) | 330 ± 44 | <0.0001** |
| Treatment (crushed rock) | -39 ± 33 | 0.25 |
| ***(c)*** *Electrical Conductivity (10-30 cm)* | | |
| Intercept | 308 ± 46 | <0.0001** |
| Site (Merced-2) | 840 ± 50 | <0.0001** |
| Site (Yolo) | -133 ± 59 | 0.06* |
| Treatment (crushed rock) | -22 ± 44 | 0.62 |
| ***(d)*** *Gravimetric Soil Water Content (10-30 cm)* | | |
| Intercept | 0.2 ± 0.01 | <0.0001** |
| Site (Merced-2) | 0.05 ± 0.01 | <0.0007** |
| Site (Yolo) | -0.05 ± 0.01 | <0.0009** |
| Treatment (crushed rock) | -0.005 ± 0.01 | 0.65 |
| **(e)** *Water Holding Capacity (0-10 cm)* | | |
| Intercept | 29.3 ± 0.6 | <0.0001** |
| Site (Merced-2) | -1.3 ± 0.75 | 0.05* |
| Site (Yolo) | -0.4 ± 0.9 | 0.55 |
| Treatment (crushed rock) | -0.8 ± 0.7 | 0.2 |
| **(f)** *Active microbial biomass (0-10 cm)* | | |
| Intercept | 21 ± 0.8 | <0.0001** |
| Site (Merced-2) | 5 ± 0.97 | 0.0001** |
| Site (Yolo) | -9.16 ± 1.1 | <0.0001** |
| Treatment (crushed rock) | -2.7 ± 0.9 | 0.004** |
| **(g)** *Active microbial biomass (10-30 cm)* | | |
| Intercept | 10.4 ± 0.7 | <0.0001** |
| Site (Merced-2) | 2.9 ± 0.9 | 0.003** |
| Site (Yolo) | -6.8 ± 1.0 | <0.0001** |
| Treatment (crushed rock) | 1.5 ± 0.8 | 0.06* |
| **(h)** *Cumulative respiration – 60 days (0-10 cm)* | | |
| Intercept | 151 ± 64 | 0.03** |

| | | |
|---|---|---|
| Site (Merced-2) | 197 ± 76 | 0.01** |
| Site (Yolo) | 169 ± 88 | 0.07* |
| Treatment (crushed rock) | 28 ± 67 | 0.68 |
| **(i)** *Cumulative respiration – 60 days (10-30 cm)* | | |
| Intercept | 132 ± 11 | <0.0001** |
| Site (Merced-2) | -23 ± 9 | 0.02** |
| Site (Yolo) | -56 ± 14 | <0.0001** |
| Treatment (crushed rock) | -0.9 ± 8 | 0.9 |
| **(j)** *Mg (0-10 cm)* | | |
| Intercept | 4.4 ± 1.4 | 0.006** |
| Site (Merced-2) | 2.9 ± 1.7 | 0.11 |
| Site (Yolo) | 19 ± 2.0 | <0.0001** |
| Treatment (crushed rock) | -1 ± 1.5 | 0.5 |
| **(k)** *Mg (10-30 cm)* | | |
| Intercept | 3.0 ± 0.6 | 0.0001** |
| Site (Merced-2) | 9.7 ± 0.8 | <0.0001** |
| Site (Yolo) | 3.6 ± 0.9 | <0.0001** |
| Treatment (crushed rock) | -0.7 ± 0.7 | 0.3 |
| **(l)** *Ca (0-10 cm)* | | |
| Intercept | 12.4 ± 3 | 0.0002** |
| Site (Merced-2) | 11.2 ± 3 | 0.003** |
| Site (Yolo) | 7.7 ± 4 | 0.06* |
| Treatment (crushed rock) | -1.0 ± 3 | 0.74 |
| **(m)** *Ca (10-30 cm)* | | |
| Intercept | 9.7 ± 2 | 0.0001** |
| Site (Merced-2) | 34.3 ± 2 | <0.0001** |
| Site (Yolo) | -4.8 ± 3 | 0.11 |
| Treatment (crushed rock) | -2.1 ± 2 | 0.34 |
| **(n)** *Na (0-10 cm)* | | |
| Intercept | 27.1 ± 4 | <0.0001** |
| Site (Merced-2) | 19.2 ± 5 | 0.0008** |
| Site (Yolo) | -19.6 ± 6 | 0.003** |
| Treatment (crushed rock) | 0.7 ± 4 | 0.9 |
| **(o)** *Na (10-30 cm)* | | |
| Intercept | 21.8 ± 1 | <0.0001** |
| Site (Merced-2) | 31.6 ± 2 | <0.0001** |
| Site (Yolo) | -18.1 ± 2 | <0.0001** |
| Treatment (crushed rock) | -0.8 ± 1 | 0.5 |
| **(p)** *Si (0-10 cm)* | | |
| Intercept | 9.2 ± 1 | <0.0001** |
| Site (Merced-2) | 2.0 ± 1 | 0.1 |
| Site (Yolo) | -1.9 ± 1 | 0..2 |
| Treatment (crushed rock) | -0.7 ± 1 | 0.5 |
| **(q)** *Si (10-30 cm)* | | |

| | | |
|---|---|---|
| Intercept | 9.0 ± 10.8 | <0.0001** |
| Site (Merced-2) | -0.6 ± 0.95 | 0.5 |
| Site (Yolo) | -0.95 ± 1 | 0.4 |
| Treatment (crushed rock) | -0.51 ± 0.8 | 0.5 |
| **(r)** *K (0-10 cm)* | | |
| Intercept | 2.7 ± 0.9 | 0.03** |
| Site (Merced-2) | 5.7 ± 0.9 | <0.0001** |
| Site (Yolo) | 21.8 ± 1.2 | <0.0001** |
| Treatment (crushed rock) | -0.1 ± 0.9 | 0.9 |
| **(s)** *K (10-30 cm)* | | |
| Intercept | 1.3 ± 0.4 | 0.002** |
| Site (Merced-2) | 3.0 ± 0.5 | <0.0001** |
| Site (Yolo) | 5.0 ± 0.5 | <0.0001** |
| Treatment (crushed rock) | 0.02 ± 0.43 | 0.97 |
| **(t)** *Fe (0-10 cm)* | | |
| Intercept | 0.3 ± 0.1 | 0.03** |
| Site (Merced-2) | -0.3 ± 0.1 | 0.05* |
| Site (Yolo) | -0.3 ± 0.2 | 0.1 |
| Treatment (crushed rock) | 0.06 ± 0.1 | 0.6 |
| **(u)** *Fe (10-30 cm)* | | |
| Intercept | 0.3 ± 0.4 | 0.3 |
| Site (Merced-2) | -0.2 ± 0.4 | 0.6 |
| Site (Yolo) | 1.4 ± 0.5 | 0.01* |
| Treatment (crushed rock) | -0.21 ± 0.4 | 0.6 |
| **(v)** *Al (0-10 cm)* | | |
| Intercept | 0.3 ± 0.1 | 0.03* |
| Site (Merced-2) | -0.3 ± 0.1 | 0.05* |
| Site (Yolo) | -0.3 ± 0.2 | 0.06* |
| Treatment (crushed rock) | 0.06 ± 0.1 | 0.6 |
| **(w)** *Al (10-30 cm)* | | |
| Intercept | 0.3 ± 0.3 | 0.3 |
| Site (Merced-2) | -0.3 ± 0.4 | 0.5 |
| Site (Yolo) | 1 ± 0.5 | 0.01** |
| Treatment (crushed rock) | -0.14 ± 0.3 | 0.6 |
| **(x)** *P (0-10 cm)* | | |
| Intercept | 2.5 ± 2 | 0.2 |
| Site (Merced-2) | 4.8 ± 2 | 0.06* |
| Site (Yolo) | -0.6 ± 3 | 0.8 |
| Treatment (crushed rock) | -2.5 ± 2 | 0.3 |
| **(y)** *P (10-30 cm)* | | |
| Intercept | 0.3 ± 0.08 | 0.002** |
| Site (Merced-2) | 0.3 ± 0.1 | 0.006** |
| Site (Yolo) | -0.2 ± 0.1 | 0.1 |
| Treatment (crushed rock) | 0.1 ± 0.09 | 0.2 |

**Table S11. Permutational analysis of variance (PERMANOVA) for bacteria and fungi at 0-10 and 10-30 cm depths respectively.** This dataset includes the three sites where sequencing data was available (Yolo, Merced-1 and Merced-2). *($p < 0.1$ is considered marginally significant and indicated by a single asterisk *; $p < 0.05$ is considered significant and indicated by two asterisks **).*

| | **Bray-Curtis** | | | **Euclidean** | | |
|---|---|---|---|---|---|---|
| **Bacteria** | **F value** | **DF** | **p value** | **F value** | **DF** | **p value** |
| 0-10 cm | | | | | | |
| Treatment | 1.3446 | 1 | 0.2392 | 0.9354 | 1 | 0.4172 |
| Site | 20.7055 | 2 | 0.0001** | 10.3609 | 2 | 0.0001** |
| Treatment*Site | 2.2127 | 2 | 0.0263** | 0.98 | 2 | 0.4389 |
| 10-30 cm | | | | | | |
| Treatment | 0.6423 | 1 | 0.7057 | 0.9354 | 1 | 0.4157 |
| Site | 11.8923 | 2 | 0.0001** | 10.3609 | 2 | 0.0001** |
| Treatment*Site | 0.7438 | 2 | 0.7039 | 0.98 | 2 | 0.4392 |
| **Fungi** | | | | | | |
| 0-10 cm | | | | | | |
| Treatment | 1.6607 | 1 | 0.0965* | 1.128 | 1 | 0.2742 |
| Site | 9.8799 | 2 | 0.0001** | 8.2797 | 2 | 0.0001** |
| Treatment*Site | 1.2437 | 2 | 0.2219 | 1.1724 | 2 | 0.2303 |

| | | | | | | |
|---|---|---|---|---|---|---|
| 10-30 cm | | | | | | |
| Treatment | 0.9151 | 1 | 0.4738 | 0.9403 | 1 | 0.4342 |
| Site | 9.6585 | 2 | 0.0001** | 6.5875 | 2 | 0.0001** |
| Treatment*Site | 1.0965 | 2 | 0.3251 | 1.0486 | 2 | 0.3392 |

**Table S12. Differential abundance analysis of unfiltered sequence data at the ASV level for bacteria and fungi at 0-10 and 10-30 cm depths respectively.** This dataset includes the three sites where sequencing data was available (Yolo, Merced-1 and Merced-2). Taxa showing significant differences between treatment and control ($p < 0.05$) across all three sites shown.

| **Data** | **Depth (cm)** | **ASV** | **Kingdom** | **Phylum** | **Class** | **Order** | **Family** | **Genus** | **Species** | **log2 Fold Change** | **p-value (adjusted)** |
|---|---|---|---|---|---|---|---|---|---|---|---|
| 16S | 0_10 | ASV_69 | Bacteria | Chloroflexi | KD4-96 | <NA> | <NA> | <NA> | <NA> | 20.1 | 0.0103 |
| 16S | 0_10 | ASV_186 | Bacteria | Actinobacteriota | Rubrobacteria | Rubrobacterales | Rubrobacteriaceae | Rubrobacter | <NA> | -4.8 | 0.0056 |
| 16S | 0_10 | ASV_645 | Bacteria | Planctomycetota | Planctomycetes | Planctomycetales | <NA> | <NA> | <NA> | 4.0 | 0.0386 |
| 16S | 0_10 | ASV_731 | Bacteria | Proteobacteria | Alphaproteobacteria | Sphingomonadales | Sphingomonadaceae | Ellin6055 | <NA> | 2.2 | 0.0616 |
| 16S | 10_30 | ASV_161 | Bacteria | Proteobacteria | Alphaproteobacteria | Sphingomonadales | Sphingomonadaceae | Sphingomonas | S. jaspsi | 4.2 | 0.0002 |
| 16S | 10_30 | ASV_299 | Bacteria | Proteobacteria | Alphaproteobacteria | Sphingomonadales | Sphingomonadaceae | Sphingomonas | S. flava | 4.9 | 0.0070 |

| | | | | | | | | | | | |
|---|---|---|---|---|---|---|---|---|---|---|---|
| 16S | 10_30 | ASV_711 | Bacteria | Verrucomicrobiota | Verrucomicrobiae | Chthoniobacterales | Chthoniobacteraceae | Candidatus | C. udaeobacter | 3.9 | 0.0561 |
| 16S | 10_30 | ASV_723 | Archaea | Crenarchaeota | Nitrososphaeria | Nitrososphaerales | Nitrososphaeraceae | <NA> | <NA> | 4.1 | 0.0443 |
| 16S | 10_30 | ASV_849 | Bacteria | Proteobacteria | Alphaproteobacteria | Sphingomonadales | Sphingomonadaceae | Ellin6055 | <NA> | 4.4 | 0.0628 |
| 16S | 10_30 | ASV_906 | Bacteria | Actinobacteriota | Thermoleophilia | Solirubrobacterales | 67-14 | <NA> | <NA> | 4.4 | 0.0628 |
| ITS | 0_10 | ASV_32 | Fungi | Ascomycota | Dothideomycetes | Pleosporales | Sporormiaceae | <NA> | <NA> | -18.2 | 0.0003 |
| ITS | 0_10 | ASV_52 | Fungi | Ascomycota | Sordariomycetes | Sordariales | Lasiosphaeriaceae | Schizothecium | S. inaequale | -18.5 | 0.0003 |
| ITS | 0_10 | ASV_68 | Fungi | Ascomycota | Sordariomycetes | Microascales | Microascaceae | Pithoascus | P. nidicola | -20.9 | 0.0005 |

| | | | | | | | | | | | |
|---|---|---|---|---|---|---|---|---|---|---|---|
| ITS | 0_10 | ASV_84 | Fungi | Ascomycota | Sordariomycetes | Microascales | Microascaceae | <NA> | <NA> | 22.5 | 0.0003 |
| ITS | 0_10 | ASV_230 | Fungi | Ascomycota | Dothideomycetes | Pleosporales | Didymellaceae | <NA> | <NA> | -18.4 | 0.0003 |
| ITS | 0_10 | ASV_429 | Fungi | Ascomycota | Pezizomycetes | Pezizales | Pezizaceae | Iodophanus | <NA> | 22.6 | 0.0004 |
| ITS | 10_30 | ASV_9 | Fungi | Ascomycota | Eurotiomycetes | Eurotiales | Aspergillaceae | Aspergillus | <NA> | -7.6 | 0.0137 |
| ITS | 10_30 | ASV_33 | Fungi | Ascomycota | Sordariomycetes | Microascales | Microascaceae | Pithoascus | P. nidicola | 22.2 | 0.0007 |
| ITS | 10_30 | ASV_52 | Fungi | Ascomycota | Sordariomycetes | Sordariales | Lasiosphaeriaceae | Schizothecium | S. inaequale | -18.6 | 0.0021 |
| ITS | 10_30 | ASV_93 | Fungi | Mortierellomycota | Mortierellomycetes | Mortierellales | Mortierellaceae | Mortierella | M. elongata | -18.4 | 0.0000 |

| | | | | | | | | | | | |
|---|---|---|---|---|---|---|---|---|---|---|---|
| ITS | 10_30 | ASV_159 | Fungi | Ascomycota | Eurotiomycetes | Onygenales | Onygenales_fam_Incertae_sedis | Chrysosporium | C. lobatum | -19.1 | 0.0004 |
| ITS | 10_30 | ASV_239 | Fungi | Basidiomycota | Agaricomycetes | Cantharellales | Ceratobasidiaceae | Thanatephorus | T. cucumeris | -22.2 | 0.0007 |
| ITS | 10_30 | ASV_277 | Fungi | Ascomycota | Sordariomycetes | Hypocreales | Nectriaceae | Gibberella | G. tricincta | 20.5 | 0.0026 |
| ITS | 10_30 | ASV_365 | Fungi | Ascomycota | Pezizomycetes | Pezizales | Pezizaceae | Iodophanus | <NA> | -23.1 | 0.0004 |
| ITS | 10_30 | ASV_545 | Fungi | Basidiomycota | Agaricomycetes | Agaricales | Entolomataceae | Entoloma | E. undatum | 17.3 | 0.0356 |
